# Translational control of breast cancer plasticity

**DOI:** 10.1101/596544

**Authors:** Michael Jewer, Laura Lee, Guihua Zhang, Jiahui Liu, Scott D. Findlay, Krista M. Vincent, Kristofferson Tandoc, Dylan Dieters-Castator, Daniela F. Quail, Indrani Dutta, Mackenzie Coatham, Zhihua Xu, Bo-Jhih Guan, Maria Hatzoglou, Andrea Brumwell, James Uniacke, Christos Patsis, Antonis Koromilas, Julia Schueler, Gabrielle M. Siegers, Ivan Topisirovic, Lynne-Marie Postovit

**Affiliations:** Department of Anatomy and Cell Biology, University of Western Ontario, London, ON, Canada; Department of Oncology, University of Alberta, Edmonton AB, Canada; Lady Davis Institute, Department of Oncology, Division of Experimental Medicine, McGill University, Montreal, QC, Canada; Goodman Cancer Center, McGill University, Montreal, QC, Canada; Department of Pharmacology, Case Western Reserve, Cleveland OH, USA; Department of Molecular and Cellular Biology, University of Guelph, Guelph, ON, Canada; Charles River Discovery Research Services Germany, Freiburg, Germany

**Author notes:** Correspondence and material requests should be directed to Dr. Lynne-Marie Postovit.

## Abstract

Plasticity of neoplasia, whereby cancer cells attain stem-cell-like properties, is required for disease progression and represents a major therapeutic challenge. We report that in breast cancer cells *NANOG*, *SNAIL* and *NODAL* transcripts manifest multiple isoforms characterized by different 5’ Untranslated Regions (5’UTRs), whereby translation of a subset of these isoforms is stimulated under hypoxia. This leads to accumulation of corresponding proteins which induce plasticity and “fate-switching” toward stem-cell like phenotypes. Surprisingly, we observed that mTOR inhibitors and chemotherapeutics induce translational activation of a subset of *NANOG*, *SNAIL* and *NODAL* mRNA isoforms akin to hypoxia, engendering stem cell-like phenotypes. Strikingly, these effects can be overcome with drugs that antagonize translational reprogramming caused by eIF2α phosphorylation (e.g. ISRIB). Collectively, our findings unravel a hitherto unappreciated mechanism of induction of plasticity of breast cancer cells, and provide a molecular basis for therapeutic strategies aimed at overcoming drug resistance and abrogating metastasis.

## Introduction

More than 20% of breast cancer patients will die due to therapy resistance and metastasis. Both processes require cancer cells to adapt to stresses like hypoxia or chemotherapy. This adaptability is thought to be mediated by increased plasticity of breast cancer cells concomitant with the presence of breast cancer cells with stem cell-like features (BCSC)^1, 2^.

Breast cancer cell plasticity and stem cell-like phenotypes are induced by hypoxia ^3–5^. Although the mechanisms by which this occurs are poorly understood, it has been reported that hypoxia induces levels of “stemness” factors including NANOG, SNAIL and NODAL ^6–8^. Moreover, both stem-cell like phenotypes and elevated expression of “stemness” factors have been linked to the metastatic spread of disease and chemoresistance ^1^. In response to hypoxia, global protein synthesis is reduced to decrease oxygen consumption. Hypoxia-induced suppression of global protein synthesis occurs via two major signaling events: Inhibition of the mammalian/mechanistic Target of Rapamycin Complex 1 (mTORC1) and induction of the Integrated Stress Response (ISR) arm of the Unfolded Protein Response (UPR) ^9–11^. MTORC1 inhibition reduces translation in part via 4E-BP-dependent inhibition of eIF4F complex assembly (eIF4E:eIF4G:eIF4A), which recruits mRNA to the ribosome ^12^. Phosphorylation of eIF2α and a decrease in ternary complex (eIF2:tRNAiMet:GTP) recycling limit initiator tRNA delivery and signify ISR ^13^. In spite of decreased global protein synthesis, a subset of mRNAs is preferentially translated during ISR, which is largely predetermined by 5’ Untranslated Region (5’UTR) features, including inhibitory upstream open reading frames (uORFs) ^14^. The inhibition of mRNA translation, mediated by mTOR suppression and eIF2α phosphorylation, drives an adaptive stress response by conserving energy whilst enabling the accumulation of adaptive factors, including ATF4 ^15^. Indeed, emerging data suggest that mTOR inhibition and eIF2α phosphorylation promote stem-cell like phenotypes. For instance, in Drosophila germline stem cells, differentiation and growth are reduced when mTOR is blocked ^16^. Furthermore, by decreasing 4E-BP phosphorylation, mTOR inhibition promotes self-renewal in neural stem cells ^17, 18^ and the phosphorylation of eIF2α is essential for the maintenance of self-renewal in satellite cells and hESCs ^19, 20^. Collectively, these studies show that translational reprograming regulates stem cell-associated plasticity, but the role of this process in the acquisition of BCSC phenotypes remains elusive.

Herein, we demonstrate that in breast cancer cells *NANOG*, *SNAIL* and *NODAL* exist in multiple isoforms which differ in their 5’UTRs, a subset of which is selectively translated under hypoxia leading to accumulation of corresponding proteins. This induction of an adaptive stem cell program leads to the acquisition of BCSC phenotypes. Strikingly this translation-regulated mechanism of plasticity is also induced by mTOR inhibitors and chemotherapeutics. Finally, we show that inhibiting the ISR with the Integrated Stress Response Inhibitor (ISRIB) impedes acquisition of stem cell-like phenotypes and therapy resistance of breast cancer cells.

## Results

### NODAL mediates the effects of hypoxia on breast cancer cell plasticity and stem cell-associated phenotypes

Stem cell associated proteins, including NANOG, SNAIL and NODAL have been shown to induce epithelial to mesenchymal transition (EMT) and are associated with poor outcomes in breast cancer patients ^21–24^. NANOG and SNAIL can also induce BCSCs ^21, 22, 25^; but notwithstanding a single report ^26^ the function of NODAL in induction of stem-like phenotypes in breast cancer cells remains unestablished. Since NODAL may co-operate with and/or induce factors such as SNAIL and NANOG to support plasticity and induction of stem-like properties ^27^, we first sought to investigate whether NODAL induces BCSC phenotypes. To this end, we employed a single cell sphere formation assay ^28, 29^. and monitored for the population of CD44high/CD24low cells ^2, 30^, which are widely used functional readouts for BCSCs. Recombinant human NODAL (rhNODAL) increased sphere formation in T47D and MCF7 cells (Figure 1a; Supplemental Figure 1a), and enriched the population of CD44high/CD24low cells in T47D, MCF7 and SUM149 cells (Supplemental Figure 1b-d). Furthermore, NODAL was up-regulated in BCSC-enriched 3D cultures (Figure 1b), and NODAL depletion or treatment with receptor kinase inhibitor that blocks NODAL signaling ^31^ (SB431542, 1μM) reduced sphere formation by MDA-MB-231 cells (Figure 1c,d). These data corroborated that NODAL promotes BCSC phenotypes.

**Figure 1:**
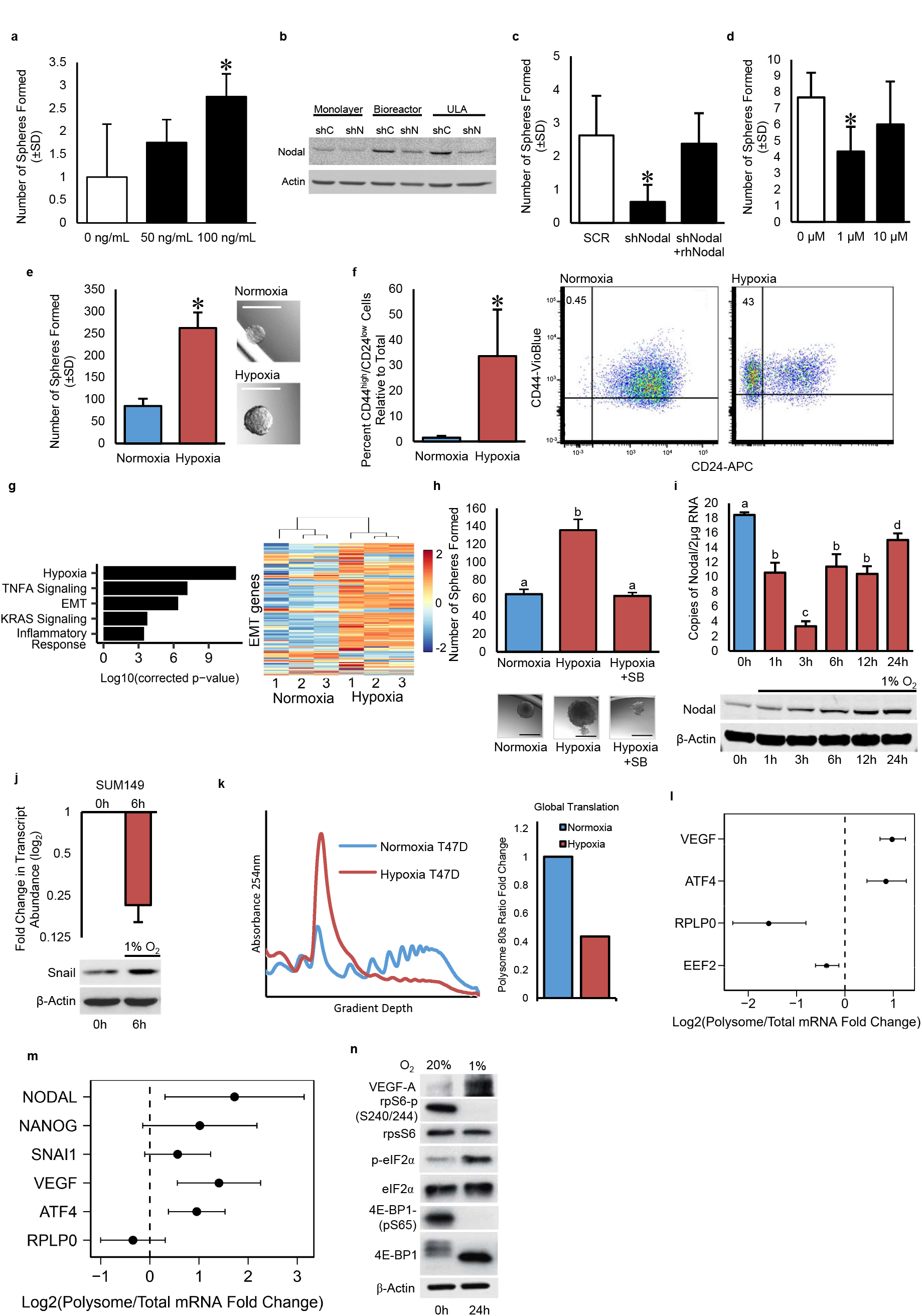
Hypoxia induces BCSC phenotypes concomitant with translational reprogramming: **a)** The number of spheres formed from T47D cells treated with rhNODAL (10, 100 ng/mL). Bars represent the mean tumorsphere count relative to untreated cells ± SD. **b)** Western blots of NODAL protein in MDA-MB-231 cells cultured in 2D or in BCSC-enriching 3D conditions (growth in a Bioreactor or Ultra Low Attachment (ULA) plates). shRNA against NODAL is used to show specificity of antibody and Actin is used as a loading control. **c,d)** The number of spheres formed from MDA-MB-231 cells expressing an shRNA against NODAL and then treated with or without rhNODAL (100 ng/mL) or **d)** exposed to SB431542 (1-10 μM). Bars represent the mean tumorsphere count relative to untreated cells ± SD. **e)** The number of spheres formed from 960 viable T47D cells pre-exposed to hypoxia or normoxia for 24h and then cultured as single cell suspensions. Bars represent the mean tumorsphere count ± SD. Images of representative spheres are presented. Micron bars = 100 μm. **f)** Percentage of T47D cells expressing CD44high/CD24low following exposure to hypoxia or normoxia for 24h. Bars represent mean percentage of CD44high/CD24low cells ± SD. Representative scatter plots discriminating subpopulations as defined by cell surface markers CD44 and CD24 are shown. **g)** Gene ontology (GO) enrichment analysis of RNAseq data from T47D cells cultured for 48h in normoxia or hypoxia confirms that hypoxia alters transcripts involved in processes critical for breast cancer plasticity. Heat map representing hierarchial clustering shows that hypoxia up-regulates genes associated with EMT. **h)** The number of spheres formed from 960 viable T47D cells pre-exposed to hypoxia or normoxia for 24h +/- SB431542 (10 μM) and then cultured as single cell suspensions. Bars represent the mean tumorsphere count ± SD. Images of representative spheres are presented. Micron bars = 100 μm. **i)** *NODAL* transcript (detected with digital RT-PCR) and protein levels (detected with Western blotting) in T47D cells cultured in hypoxia for 0-24h. Bars represent absolute copies of NODAL and Actin is used as loading control for Western blotting. Different letters are statistically different. **j)** *SNAIL* transcript (detected with real time RT-PCR) and protein levels (detected with Western blotting) in SUM149 cells cultured in hypoxia for 0 and 6h. Bars represent Log2 fold change in transcript relative to levels at 0h and Actin is used as loading control for Western blotting. **k)** Translation rates in cells used in a) represented as fold change of mRNA associated with polysomes (more than 3 ribosomes) in cells cultured in hypoxia versus normoxia. **l,m)** Polysome associated *EEF2*, *RPLPO*, *VEGF* and *ATF4* levels in T47D cells (**l**) and *RPLPO*, *VEGF*, *ATF4*, *NANOG*, *SNAIL* and *NODAL* levels in H9 hESC (**m**) cultured for 24h in 1 or 20% O_2_. Levels in polysomes were normalized to total mRNA levels and graph represents the mean Log2 fold change in hypoxia relative to levels in normoxia ± SD. **n)** Western blots of lysates from T47D breast cancer cells exposed to 20 or 1% O_2_ for 24h show that low O_2_ reduces 4E-BP and rpS6 phosphorylation, and increases eIF2α phosphorylation. Total 4E-BP, rpS6 and eIF2α levels were unchanged. B-Actin is used as a loading control and VEGF is used as a positive control for the hypoxic response. Polysome profiles of lysates extracted from the same cells are presented. Unless indicated otherwise, for all graphs, asterisks (*) indicate a significant difference from control and different letters are statistically different from each other (p<0.05).

Hypoxia is thought to act as a major factor that increases plasticity of cancer cells leading to the establishment of BCSC phenotypes. We therefore next sought to determine the role of stem cell factors (e.g. NANOG, SNAIL and NODAL) in mediating hypoxia-induced BCSC phenotypes. T47D and SUM149 cells were maintained under hypoxia for 24 hours. Although hypoxia did not alter viability during this time course, we observed hypoxia-associated increases in tumorsphere formation, as well as the emergence of a larger CD44high/CD24low population (Figure 1e,f; Supplemental Figure 1e). As this dose of hypoxia was not lethal (and thus did not selectively cause cell death in non-BCSC populations), these findings suggest that under hypoxia BCSCs are induced, rather than selected for. Moreover, RNA sequencing of T47D cells cultured for 48 hours in 20 or 1% O_2_ demonstrated that hypoxia also entices an EMT signature (Figure 1g; Supplemental Table 1). Finally, SB431542 treatment abrogated the hypoxic induction of sphere formation in T47D and SUM149 cells, further corroborating that NODAL facilitates plasticity in response to hypoxia (Figure 1h; Supplemental Figure 1f).

### Hypoxia alters the levels of NODAL, NANOG and SNAIL by modulating translation of corresponding mRNAs

To investigate the underlying mechanisms of the observed role of NODAL in increasing breast cancer cell plasticity and engendering stem cell-like phenotypes in breast cancer cells, we next examined NODAL protein and mRNA levels in T47D breast cancer cells exposed to hypoxia for 6-24 hours. Unexpectedly, we observed that although NODAL protein was up-regulated after 3-24 hours of hypoxia (1% O_2_), the levels of corresponding mRNA were downregulated after 3 hours followed by a partial recovery after 24 hours (Figure 1i). Strikingly, similar discordance between mRNA and protein levels was also observed for SNAIL and NANOG (Figure 1j)(Supplemental Figure 1g). These findings strongly suggest that the levels of NODAL, SNAIL and NANOG proteins under hypoxia are regulated at the level of translation.

We next employed polysome profiling, which separates efficiently versus non-efficiently translated mRNAs by ultracentrifugation on sucrose gradients ^32^. This confirmed that exposure to hypoxia for 24 hours causes a dramatic (70-95%) reduction in global translation in T47D, MCF7, and H9 cells (Figure 1k, Supplemental Figure 1h,i). Using digital droplet RT-PCR (ddPCR) on total and efficiently translated mRNA fractions (associated with more than 3 ribosomes), we next assessed the effects of hypoxia on mRNAs representative of known 5’UTR-dependent modes of translational regulation ^14^. As expected, in T47D cells hypoxia reduced translation of 5′ terminal oligopyrimidine (TOP) mRNAs Eukaryotic Elongation Factor 2 (*EEF2*) and Ribosomal Protein, Large, P0 (*RPLPO*) (Figure 1l) while enhancing the translation of Activating Transcription Factor 4 (*ATF4*) and Vascular Endothelial Growth Factor (*VEGF*) mRNAs (Figure 1l). Strikingly, using hESCs, we noted that similarly to *VEGF* and *ATF4* and in contrast to *RPLPO*, the translation of *NANOG*, *NODAL* and *SNAIL* mRNA was either sustained or increased under hypoxia (Figure 1m). Stresses such as hypoxia cause adaptive translational reprogramming via modulating mTOR and integrated stress response (ISR) signaling pathways ^33–36^. Using Western blotting, we confirmed that in T47D cells hypoxia reduces mTORC1 activity - as illustrated by reduced 4E-BP1 and ribosomal protein S6 (rpS6) phosphorylation (1% O_2_; 24 hours) - while inducing ISR as evidenced by eIF2α phosphorylation (Figure 1 n; Supplemental Figure 1j, 1% O_2_). In T47D and MCF7 breast cancer cells as well as H9 human embryonic stem cells (hESC), we further demonstrated that, depending on the cell type, hypoxia-induced perturbations in mTOR activity and eIF2α phosphorylation are initiated between 3 and 12 hours (Supplemental Figure 1j). Collectively, these data confirm the previous findings that hypoxia induces translational reprogramming by inhibiting mTORC1 and bolstering eIF2α phosphorylation. More importantly, these results suggest that the effects of hypoxia on translation of mRNAs encoding “stemness” factors NANOG, NODAL and SNAIL are distinct from the regulatory patterns observed for TOP mRNAs (e.g. EEF2, RPLP0), and resemble those documented for ISR-induced (e.g. ATF4) or cap-independently translated (VEGF) mRNAs.

### Multiple isoforms with distinct 5’UTR features enable translation of *SNAIL*, *NODAL* and *NANOG* mRNAs in hypoxia

Translational efficiency is determined by 5’UTR features ^14^. We therefore set out to determine the mechanism by which translation of *SNAIL*, *NODAL* and *NANOG* mRNAs is maintained under hypoxia. Using RefSeq and publicly available CAGE data, in combination with 5’RACE analysis, we made the surprising discovery that the *NODAL*, *SNAIL* and *NANOG* genes contain multiple transcriptional start sites (TSSs), which result in isoforms which differ in their 5’ UTRs (Figure 2 a-c). In the *NANOG* locus, we validated a previously described 5’UTR composed of 350 nucleotides (nt) ^37^ as well as an alternative 5’UTR of 291 nt (Figure 2a). We observed 2 TSS in the *SNAIL* locus: a distal site that yields a 5’UTR of 417 nt and a proximal site, which generates a 5’UTRof 85 nt (Figure 2b). In the *NODAL* locus there were four 5’UTRs comprised of 14, 42 and 298 and 416 nt (Figure 2c). We next investigated whether NODAL, SNAIL and NANOG isoforms are differentially translated under hypoxia. MCF7 cells were cultured in 20 or 1% O_2_ for 24 hours, and then fractionated to obtain mRNAs associated with monosomes, light (less than 3 ribosomes) and heavy (more than 3 ribosomes) polysomes (Supplemental Figure 2a). The percentage of transcript associated with each fraction was then determined using isoform-specific quantitative RT-PCR. As a control, we confirmed that *ATF4* mRNA was shifted towards heavy polysomes in cells cultured in hypoxia versus normoxia (Figure 2d). Parallel results were observed for *VEGFA* mRNA, which is also selectively translated in hypoxia ^38^ (Supplemental Figure 2b). This is in contrast to *ACTIN*, which exhibited an expected hypoxia-associated reduction in translational efficiency (Supplemental Figure 2c). Using this method, we showed that the shorter NANOG 291 mRNA exerts stronger shift to heavy polysomes relative to the longer isoform (Figure 2e). In turn, the longer *SNAIL* mRNA isoform was less efficiently translated in hypoxia as compared to the shorter 5’UTR isoform (Figure 2f). Finally, translational efficiency of the *NODAL* 298 5’UTR mRNA, but not of short NODAL isoforms, was increased under hypoxia (Figure 2g). These findings suggest a previously unprecedented model of translational regulation whereby differences in 5’UTR features of the mRNA isoforms harboring the same ORF allow isoform-specific translation and consequent upregulation of NANOG, SNAIL and NODAL proteins under hypoxia.

**Figure 2:**
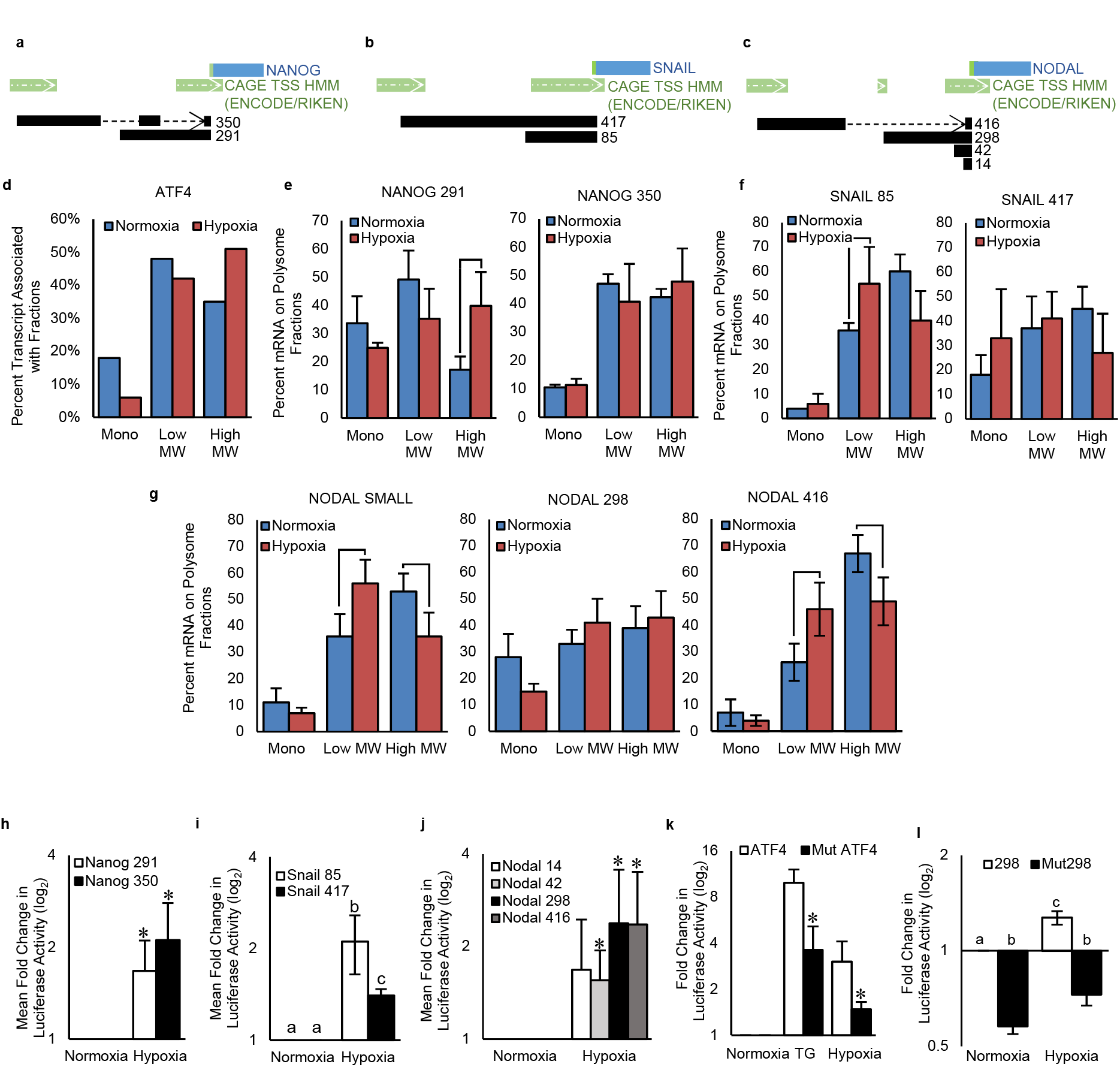
Hypoxia induces the selective translation of BCSC-associated transcripts in an isoform-specific manner. **a-c)** Diagrams of the **a)** NANOG, **b)** SNAIL and **c)** NODAL gene loci showing multiple transcriptional start sites (dark green with white arrows) leading to multiple 5’UTR sequences (black). Dotted lines represent splicing events. The blue rectangles represent the ORFs and the bright green bar represents the translational start site. **d)** *ATF4* mRNA associated with mononsomes, low MW polysomes and high MW polysomes extracted from MCF7 cells cultured for 24h in hypoxia or normoxia. Bars represent the percent of transcript associated with each fraction in each condition. A representative biological replicate is shown. **e-g)** mRNA isoforms of **e)** *NANOG*, **f)** *SNAIL* and **g)** *NODAL* associated from mononsomes, low MW polysomes and high MW polysomes extracted from MCF7 cells cultured for 24h in hypoxia or normoxia. Bars represent the mean percent of transcript associated with each fraction in each condition ± SD (n=3). Lines indicate significant differences between conditions (p<0.05). **h-j)** Luciferase activity in cells transfected with p5’UTR firefly constructs harboring the various 5’UTRs of **h)** NANOG (n=3), **i)** SNAIL (n=3), or **j)** NODAL (n=4) and then exposed for 6h to normoxia, or hypoxia. Bars represent mean luciferase activity in hypoxia relative to normoxia ± SD. Different letters are statistically different in h) and values indicated by an asterisk (*) in i) and j) are statistically different from control values but not from each other (p<0.05). **k)** Luciferase activity in cells transfected with p5’UTR firefly constructs harboring either the wild type or uORF1 mutated ATF4 5’UTR and then exposed for 6h to normoxia, thapsigargin (TG; 0.1 μM), or hypoxia. Bars represent mean luciferase activity relative to cells exposed to normoxia ± SD. Values indicated by an asterisk (*) are statistically different from unmutated 5’UTR (p<0.05, n=3). **l)** Luciferase activity in cells transfected with p5’UTR firefly constructs harboring either the wild type or uORF mutated NODAL 298 5’UTR and then exposed for 6h to normoxia or hypoxia. Bars represent mean luciferase activity relative to cells exposed to normoxia ± SD (n=3). Different letters are statistically different from each other (p<0.05).

To test this model, we cloned each 5’UTR variant of NANOG, SNAIL and NODAL isoforms into a p5’UTR firefly construct ^39^, transfected cells, exposed them to 20 or 1% O_2_ and then measured luciferase activity. In accordance with the polysome profiling data, both 5’UTRs of NANOG facilitated translation in hypoxia, whereas the shorter SNAIL 5’UTR facilitated translation significantly more than the longer SNAIL 5’UTR (Figure 2 h,i). We also found that the 416 and 298 nt NODAL 5’UTRs increased translational efficiency in hypoxia more than the 42 nt or 14 nt 5’UTRs (Figure 2j). Based on sequence analysis, we noted that the 298 nt NODAL 5’UTR contains a putative uORF (Supplemental Figure 2d). We thus sought to establish the functionality of this uORF in NODAL, by comparing the effects of the NODAL 298 and the ATF4 5’UTRs on translation, whereby ATF4 is a well-established mRNA that is translationally activated during the ISR in an uORF-dependent manner ^39–41^. Cells transfected with p5’UTR firefly constructs harboring a wildtype or uORF1 mutant ATF4 5’UTR ^39^ were exposed to 20% O_2_, 1% O_2_ or thapsigargin (TG; 0.1μM as a positive control for ISR) for 6 hours followed by measurement of luciferase activity. As expected, luciferase activity in the cells expressing constructs harboring the ATF4 5’UTR was enhanced by both hypoxia and TG, and this was abrogated by the uORF1 mutation (Figure 2k). Notably, some translation occurred in the uORF1 mutant, potentially due to recently described eIF4F-independent mechanisms of ATF4 mRNA translation during stress ^15^. Similar to the ATF4 5’UTR, translation driven by the NODAL 298 5’UTR was enhanced by hypoxia and this effect was attenuated when the uORF was mutated (Figure 2l). Altogether, these findings show that NODAL expression under hypoxia is achieved through selective translation of longer isoforms which harbor uORFs in their 5’UTR. Of note, this mode of translational regulation is markedly distinct from ATF4 translational activation under ISR, wherein a single ATF4 isoform is activated via delayed re-initiation, rather than via selection of isoforms harboring a uORF as it is the case for NODAL. Moreover, and in stark contrast to NODAL, shorter 5’UTR isoforms of *SNAIL* and *NANOG* which are devoid of putative uORFs are translated more efficiently under hypoxia than their longer counterparts. This may be due to the lower energy requirements needed to translate shorter and less complex 5’UTRs. Notwithstanding differences in the mechanisms by which 5’UTR features regulate translation of *NANOG*, *SNAIL* and *NODAL* mRNAs in hypoxia, these results indicate that the selective isoform specific translational activation of *NANOG*, *SNAIL* and *NODAL* mRNAs underpins the increase in their protein levels under hypoxia.

### mTOR inhibition induces BCSC phenotypes

The results of clinical trials in which mTOR inhibitors were used to treat breast cancer patients were less efficacious than expected, with limited survival benefits^42^. Since hypoxia reduces mTOR activity (Figure 1n) and induces plasticity and stem cell-like phenotypes in breast cancer cells (Figure 1e,f,g), we next investigated the role of mTOR in induction of NANOG, SNAIL and NODAL and stem cell-like phenotypes in hypoxia. T47D and MCF7 cells were treated for 1-24 hours with the active-site mTOR inhibitor INK128 (INK; 20nM). INK128 induced NODAL protein levels in T47D and MCF7 cells within 6 hours, as compared to the control (Figure 3a; Supplemental Figure 3a). These experiments were repeated in the presence or absence of SB431542 (10 μM) to abrogate NODAL activity. Strikingly, INK128 increased sphere forming frequency and anchorage independent growth to the same extent as rhNODAL (100ng/mL), which was used as a positive control (Figure 3b,c; Supplemental Figure 3b-e). Moreover, the effects of INK128 on sphere formation and anchorage independent growth were attenuated by SB431542, suggesting that the NODAL signaling pathway mediates the effects of INK128 on BCSC phenotypes. Consistently, the percentage of CD44high/CD24low cells in T47D, MCF7 and SUM149 cell lines was increased by INK128 and that this effect was blocked with SB431542 (Figure 3d; Supplemental Figure 3f,g). BCSCs are also characterized by the expression of stem cell markers such as NANOG and SOX2, and have typically undergone EMT ^1^. INK128 induced a stem-cell-like EMT signature in T47D and SUM149 breast cancer cells relative to vehicle-treated control cells (Figure 3e,f; Supplemental Figure 3h,i). Importantly, this effect was maintained even after a 24 hour wash out period, suggesting that INK128 may induce a reprogramming event which engenders plasticity and sustains stem cell-like properties of breast cancer cells. Considering that the latter phenomena are closely linked to metastasis and to confirm the observed effects *in vivo*, we treated breast cancer cells with INK128 for 24 hours and then injected them into the lungs of mice through the tail vein. Metastases were quantified 8 weeks later using HLA staining in order to establish tumor initiating frequency *in vivo* without confounding variables (such as alterations in processes such as angiogenesis needed for larger tumor growth) that may affect the interpretation of subcutaneous limiting dilution assays. Using this method, we determined that INK128 pretreatment increases the ability of SUM149 cells to initiate metastatic tumor formation in the lung (Figure 3g).

**Figure 3:**
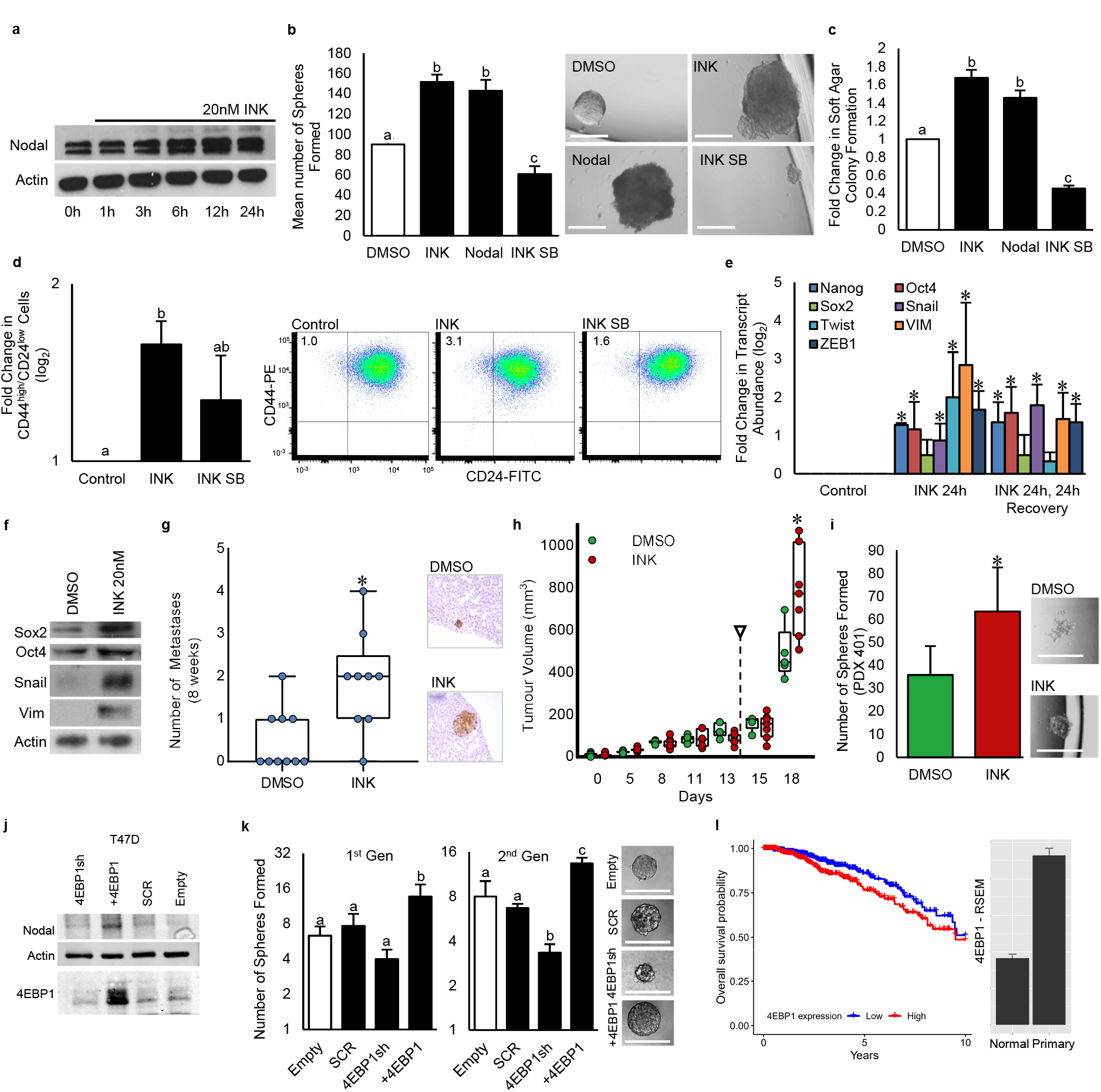
MTOR inhibition induces breast cancer plasticity: **a)** Western blot of lysates from T47D cells treated for 0-24h with MLN0128/INK128 (INK; 20nM). NODAL protein increases over time and Actin is used as a loading control. **b)** The number of spheres formed from 960 viable T47D cells pre-exposed to DMSO, INK (20nM), rhNODAL (100 ng/mL) or INK + SB431542 (10 μM) for 24h and then cultured as single cell suspensions. Bars represent the mean tumorsphere count ± SD (n=3). Images of representative spheres are presented. Micron bars = 100 μm. **c)** The number of colonies formed from T47D cells pre-exposed to DMSO, INK (20nM), rhNODAL (100 ng/mL) or INK + SB431542 (10 μM) for 24h and then cultured in soft agar. Bars represent the mean colony count ± SD relative to colonies formed from cells treated with DMSO (n=3). **d)** Percentage of T47D cells expressing CD44high/CD24low following exposure to DMSO, INK or INK+SB431542 as in b). Bars represent mean percentage of CD44high/CD24low cells ± SD (n=6). Representative scatter plots discriminating subpopulations as defined by cell surface markers CD44 and CD24 are shown. **e)** Real time RT-PCR based quantification of transcripts associated with stem cells (NANOG, SOX2, OCT4) or EMT (TWIST, ZEB1, SNAIL, VIM) in T47D cells cultured for 24h in DMSO or INK (20 nM). A group wherein INK was washed out for 24h was also included. Bars represent mean Log2 fold change relative to cells treated with DMSO ± SD (n=3). **f)** Western blot analyses of proteins associated with stem cells (SOX2, OCT4) or EMT (SNAIL, VIM) in T47D cells cultured for 24h in DMSO or INK (20 nM). Actin is used as a loading control. **g)** Lung colonies in mice 8 weeks following injection, through the tail vein, of SUM149 cells pretreated for 24h with DMSO or INK (20nM) (n=10). Representative images of cancer colonies (brown) in the lungs of mice are presented. **h)** PDX 401 tumor volumes in mice treated every second day for 2 weeks with either DMSO or INK (30 mg/kg). Arrow indicates cessation of treatment. Data presented as mean tumor volume ± SD. Time point indicated by an asterisk (*) shows significant difference between groups (p<0.01, n=8). **i)** The number of spheres formed from 96000 viable PDX 401 cells dissociated from 1cm diameter tumors taken from mice in g) and then cultured in single cell suspensions. Bars represent the mean tumorsphere count ± SD (DMSO n=6, INK n=7). Images of representative spheres are presented. Micron bars = 100 μm. **j)** Western blot analyses of NODAL and 4E-BP1 in T47D cells stably transfected with 4E-BP1 shRNA, a shRNA scrambled (SCR) control, a 4E-BP1 ORF or an empty vector control. Actin is used as a loading control. **k)** The number of spheres formed from 96 T47D cells transfected with vectors as described in j) and then cultured as single cell suspensions. Self-renewal was further assessed by measuring sphere formation in the 2^nd^ generation. Bars represent the mean tumorsphere count ± SD (n=3). Images of representative spheres are presented. Micron bars = 100 μm. **l)** Kaplan-Meier plot demonstrating the correlation between 4E-BP1 expression and survival in 1100 breast cancer patient samples analyzed by RNA-Seq. Low levels of 4E-BP1 predict survival within the first 2000 days (5.4 years, p=0.0187). Expression of the 4E-BP1 transcript increases in tumors above levels found in healthy adjacent breast tissue as determined with RNA-Seq by Expectation Maximization (RSEM) to estimate transcript abundance. For graphs, values indicated by an asterisk (*) are statistically different from controls and values indicated by letters are statistically different from each other (p<0.05).

In order to determine whether mTOR inhibition may increase populations of stem-like cells in established heterogeneous tumors, we next treated mice harboring a triple negative breast cancer patient derived xenograft (PDX 401) with INK128 (30 mg/kg, every second day for 2 weeks starting when tumor had a diameter of 5 mm). Immunohistochemical (IHC) analyses confirmed that phopsho-4E-BP1 levels were inversely correlated with hypoxia [delineated by Carbonic Anhydrase 9 (CA9)] and, as expected, were downregulated in mice treated with INK128 (Supplemental Figure 3j). Strikingly, tumor growth was not altered by INK128 during the treatment period and actually increased in INK128-treated animals after the treatment stopped (Figure 3h). Moreover, cancer cells dissociated from tumors grown in INK128-treated animals had higher BCSC frequencies than did those treated with vehicle (Figure 3i). Collectively these data provide strong evidence that mTOR inhibition may induce breast cancer cell plasticity and stemness.

We next sought to determine whether modulating the translational machinery downstream of mTOR could regulate NODAL expression. To this end, we altered the expression of 4E-BP1 that acts as a major mediator of the effects of mTOR on translation in particular in cancer ^43^ in T47D and MCF7 cells. 4E-BP1 overexpression increased NODAL levels in both cell lines (Figure 3j; Supplemental Figure 3k). NODAL levels were relatively unaffected by 4E-BP1 knock-down, likely due to low basal levels of NODAL and redundancies associated with the expression of 4E-BP2 and 4E-BP3. Importantly, 4E-BP1 overexpression also increased sphere formation (Figure 3k; Supplemental Figure 3l). Previous studies have shown the 4E-BP1 levels are high in breast tumors and that 4E-BP1 may participate in hypoxia-associated translational reprogramming ^38^. Accordingly, analysis of RNA sequencing data from 1100 breast cancer patients in the TCGA demonstrated that high levels of 4E-BP1 in the primary tumor predict poor survival in the first 5 years post-diagnosis (Figure 3l).

### The ISR mediates BCSC induction in response to stress and mTOR inhibition

Hypoxia ^11, 41^ as well as acute mTOR inhibition ^44, 45^ have been shown to enhance eIF2α phosphorylation. Accordingly, we observed that hypoxia (Figure 1n; Supplemental Figure 1j) and INK128 treatment (3-6 hours) increase eIF2α phosphorylation in T47D and SUM149 cells (Figure 4a; Supplemental Figure 4a). Thus, we next explored the extent to which the eIF2α phosphorylation may mediate BCSC induction in response to hypoxia and/or mTOR inhibition.

**Figure 4:**
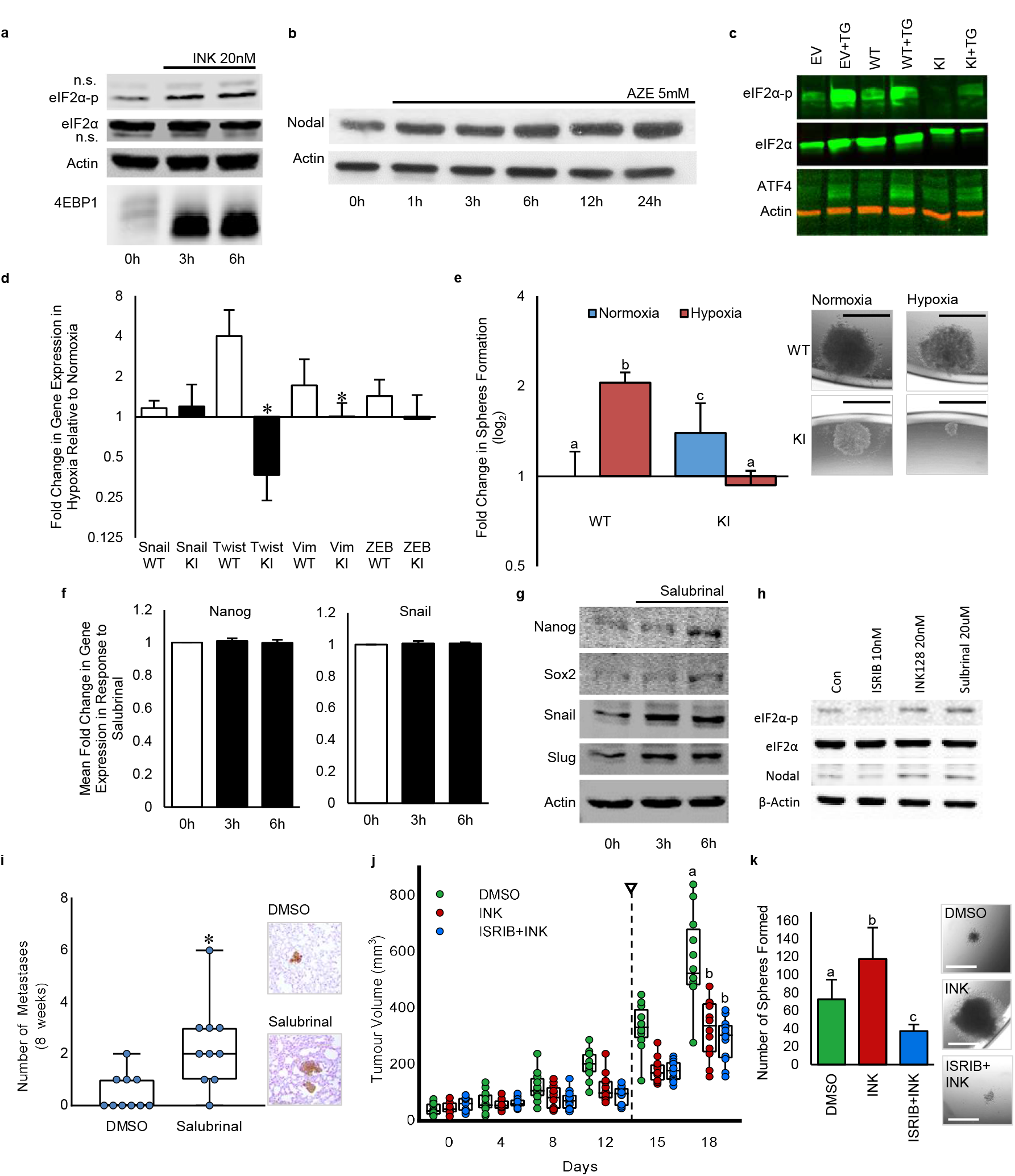
Stress-induced plasticity is mediated by the ISR: **a)** Western blots of eIF2α-p, eIF2α and 4E-BP1 in lysates extracted from T47D cells treated for 0, 3 or 6h with INK 20 nM show that INK induces eIF2α phosphorylation. Actin is used as a loading control. **b)** Western blot of NODAL in lysates from T47D cells treated with AZ (5mM) for 0-24h shows that this ER stress induces NODAL. Actin is used as a loading control. **c)** Western blots of lysates extracted from MDA-MB-231 cells expressing an empty vector (EV), a wild type eIF2α (WT) or an eIF2α mutant that cannot be phosphorylated (KI). Cells were treated with vehicle or TG. Westerns blots for eIF2α-p and ATF4 show that KI mitigates TG-induced ATF4. eIF2α and Actin are used as loading controls. **d)** Real time RT-PCR based quantification of transcripts associated with EMT (TWIST, ZEB1, SNAIL, VIM) in MDA-MB-231 cells expressing eIF2α constructs described in c) and cultured for 24h in normoxia or hypoxia (n=3). Bars represent mean Log2 fold change relative to cells cultured in normoxia ± SD. KI prevented the hypoxic induction of ZEB1, VIM and TWIST. **e)** The number of spheres formed by MDA-MB-231 variants described in c) pre-exposed to hypoxia or normoxia for 24h and then cultured as single cell suspensions. Bars represent the mean tumorsphere count relative to spheres formed by WT eIF2α expressing cells cultured in normoxia ± SD (n=3). Images of representative spheres are presented. **f)** Real time RT-PCR based quantification of NANOG and SNAIL mRNA levels in T47D cells treated for 0,3 or 6h with Salubrinal (20μM). Bars represent mean Log2 fold change relative to control (n=4). **g)** Western bot analyses of stem cell (NANOG, SOX2) and EMT (SNAIL, SLUG) proteins in T47D cells treated for 0,3 or 6h with Salubrinal (20μM). Actin is used as a loading control. **h)** Western blots of eIF2α-p and eIF2α from lysates extracted from T47D cells treated for 24h with vehicle, ISRIB (10nM), INK 20 nM or Salubrinal (20μM) show that INK and Salubrinal induce eIF2α phosphorylation and NODAL, and that ISRIB inhibits NODAL. Actin and eIF2α are used as a loading controls. **i)** Lung colonies in mice 8 weeks following injection, through the tail vein, of SUM149 cells pretreated for 24h with DMSO or Salubrinal (20μM) (n=10). Representative images of cancer colonies (brown) in the lungs of mice are presented. **j)** SUM 149 tumor volumes in mice treated every second day for 2 weeks with DMSO, INK (30 mg/kg) or INK + ISRIB (2.5 mg/kg). Arrow indicates cessation of treatment. Data are presented as mean tumor volume ± SD. Time point indicated by a letter shows significant difference between groups (p<0.01, n=7). **k)** The number of spheres formed from 9600 viable SUM 149 cells dissociated from 1cm diameter tumors taken from mice in **l)** and then cultured in single cell suspensions. Bars represent the mean tumorsphere count ± SD. Images of representative spheres are presented. Micron bars = 100 μm. For graphs, values indicated by an asterisk (*) are statistically different from controls and values indicated by letters are statistically different from each other (p<0.05).

We first activated the ISR in T47D and MCF7 cells by exposing them to azetidine-2-carboxylate (AZE; 5mM) for 1-24 hours. Using Western blotting, we determined that AZE induced NODAL within the first 6 hours of exposure (Figure 4b; Supplemental Figure 4b). Next, we depleted endogenous eIF2α in MDA-MB-231 cells and then rescued eIF2α with either a wild type (WT) or a non-phosphorylatable eIF2α mutant (S51A) ^46^. As expected, expression of the S51A eIF2α mutant prevented up-regulation of ATF4 protein in response to TG (0.1 μM; 6 hours) (Figure 4c). Moreover, exposure of MDA-MB-231 to 0.5% O2 for 24 hours increased the expression of stem cell associated genes and sphere forming frequency in cells expressing the WT but not mutant eIF2α (Figure 4d,e). This suggests that eIF2α phosphorylation is needed for the hypoxic induction of breast cancer plasticity and stem cell-like phenotypes.

To corroborate these findings, we treated breast cancer cells for 24 hours with Salubrinal (which increases phospho-eIF2α by inhibiting PP1/GADD34) and then measured the expression of stem cell and EMT markers. Salubrinal (10 μM) induced NODAL protein levels in SUM149 cells similar to those observed with INK128 treatment (Figure 3a; Supplemental Figure 3a). Using quantitative RT-PCR and Western blotting we determined that Salubrinal (up to 6 hour treatment) induced NANOG and SNAIL protein, but not mRNA, in SUM149 cells (Figure 4f,g). Conversely, NODAL levels were reduced by ISRIB (10nM), which counteracts the eIF2α-dependent inhibition of TC recycling by bolstering eIF2B GEF activity ^47, 48^ (Figure 4h). As an *in vivo* extension, we exposed breast cancer cells to Salubrinal for 24 hours and then injected them into the lungs of mice through the tail vein. We then quantified metastases 8 weeks later using HLA staining and determined that pre-exposure to Salubrinal increased the initiation of metastatic colonies in a manner similar to that observed in response to INK (Figure 4i; Figure 3g), suggesting that mTOR and the ISR may cross-talk to facilitate the emergence of breast cancer cell plasticity.

We next wanted to assess the extent to which mTOR and the ISR crosstalk during the acquisition of BCSC phenotypes. To this end, we first treated mice harboring SUM149 xenograft tumors with INK128 (30 mg/kg) or an INK128 (30 mg/kg) + ISRIB (2.5 mg/kg) in combination. Mice were dosed every second day for 2 weeks starting when tumors reached a diameter of 5mm. IHC analyses confirmed that phospho-4E-BP1 levels were inversely correlated with hypoxia (delineated by CA9) and that they were reduced in mice treated with INK128 as compared to the controls (Supplemental Figure 4c). Moreover ATF4 levels were induced by INK128, whereby ISRIB abrogated these effects (Supplemental Figure 4d). In the SUM149 xenograft model, tumor growth was reduced by both treatments (Figure 4j). Cancer cells dissociated from tumors grown in INK128-treated animals had higher BCSC frequencies than did those treated with vehicle, and this effect of INK was abolished by ISRIB (Figure 4k). Since BCSC frequencies were affected in the absence of differences in overall tumor growth, it is likely that ISRIB specifically targets the acquisition of BCSCs in this model. Collectively these data suggest that the effects of mTOR inhibition on BCSC phenotypes are mediated, at least in part, by the ISR and that ISRIB may be used to mitigate plasticity induced by hypoxia and/or TOR inhibitors, and thus improve therapeutic responses and disease outcomes.

### ISRIB abrogates chemotherapy-induced breast cancer plasticity

Chemotherapy has been shown to induce BCSCs ^49^. Paclitaxel is a first line chemotherapy used for the treatment of breast cancer. We observed that paclitaxel (20nM) treatment of T47D and SUM149 cells increases eIF2α phosphorylation while reducing phosphor-4E-BP1 levels (Figure 5a; Supplemental Figure 5a). Moreover, paclitaxel induced expression of stem cell and EMT-associated proteins including NANOG and SOX2 within 3-6 hours of treatment (Figure 5b). This confirmed that in a manner similar to hypoxia and INK128, paclitaxel induces the ISR, and suppresses mTOR signaling which coincides with acquisition of stem cell-like phenotypes.

**Figure 5:**
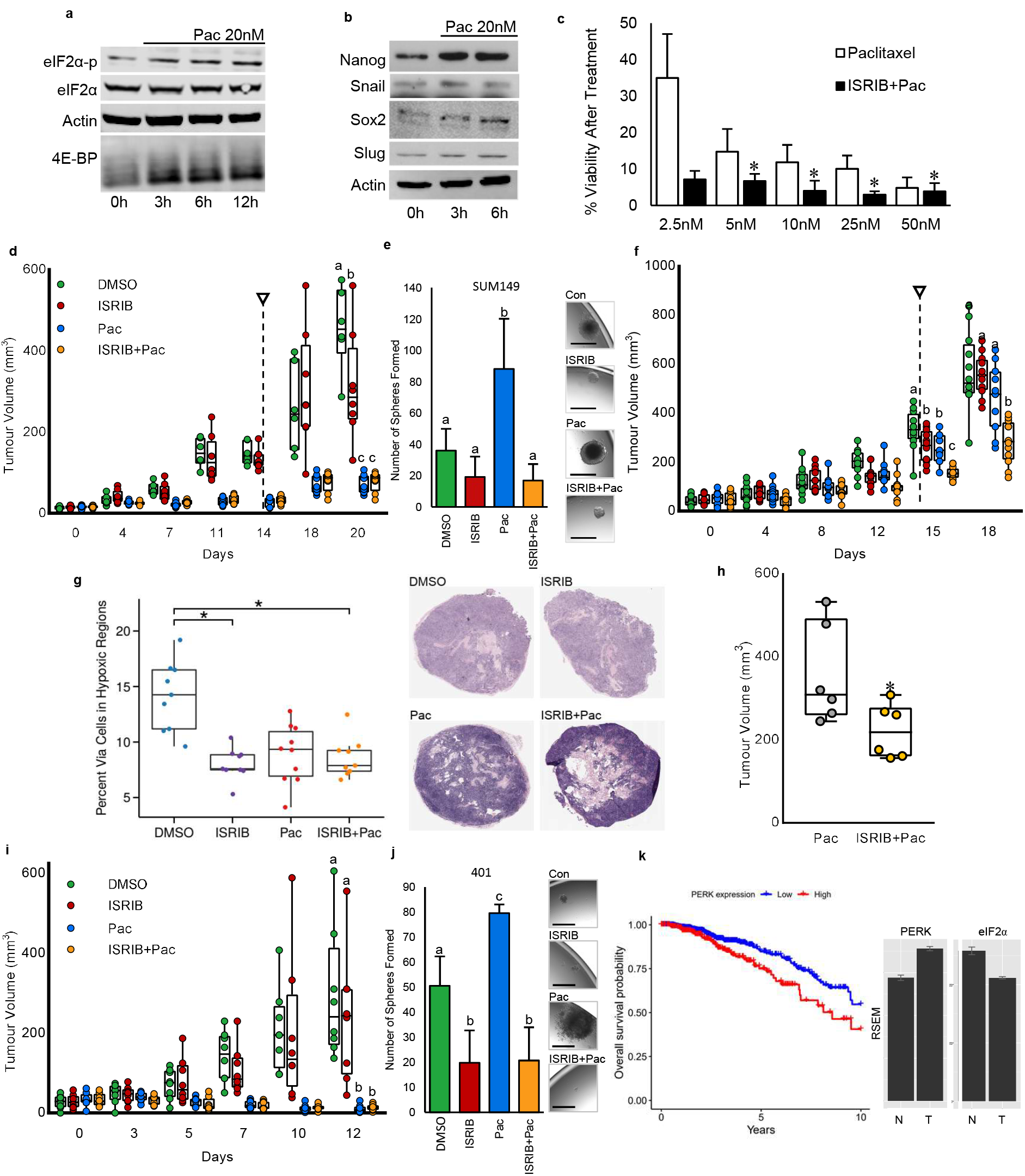
ISRIB mitigates therapy-induced BCSCs and improves efficacy of paclitaxel: **a)** Western blots of eIF2α-p, eIF2α and 4E-BP1 in lysates extracted from T47D cells treated for 0-12h with paclitaxel (20nM) show that paclitaxel induces eIF2α phosphorylation and an increase in 4E-BP1. Actin is used as a loading control. **b)** Western blots of NANOG and SOX2 in lysates extracted from T47D cells treated for 0, 3 or 6h with paclitaxel (20nM) show that paclitaxel increases NANOG and SOX2. Actin is used as a loading control. **c)** Surviving fractions of T47D cells exposed to increasing doses of paclitaxel (2.5-50 nM) in the presence or absence of ISRIB (10nM). Bars represent mean surviving fraction ± SD (n=3). Values indicated by an asterisk (*) mark doses wherein ISRIB significantly increased the efficacy of paclitaxel (p<0.05, n=3). **d)** SUM 149 tumor volumes in mice treated every second day for 2 weeks with DMSO, ISRIB (2.5 mg/kg IP), paclitaxel (15 mg/kg IV), or paclitaxel (15 mg/kg IV) + ISRIB (2.5 mg/kg IP). Arrow indicates cessation of treatment. Data presented as mean tumor volume ± SD. Time point indicated by letters shows significant difference between groups (p<0.01, treatment n=8, DMSO n=6). **e)** The number of spheres formed from 9600 viable SUM 149 cells dissociated from 1cm diameter tumors taken from mice in d) and then cultured in single cell suspensions. Bars represent the mean tumorsphere count ± SD. Images of representative spheres are presented. Micron bars = 100 μm. **f)** SUM 149 tumor volumes in mice treated every second day for 2 weeks with DMSO, ISRIB (2.5 mg/kg IP), paclitaxel (20 mg/kg IP), or paclitaxel (20 mg/kg IP) + ISRIB (2.5 mg/kg IP). Arrow indicates cessation of treatment. Data presented as mean tumor volume ± SD. Time point indicated by letters shows significant difference between groups (p<0.01, n=10). **g)** H&E staining in SUM149 tumors from f). Accompanying graph represents the mean percentage of viable cells within necrotic areas ± SD. Asterisk (*) shows significant difference (p<0.01, n=7). **h)** MDA-MB-231 tumor volumes 7 days after treating mice for 3weeks with DMSO, ISRIB (2.5 mg/kg IP), paclitaxel (15 mg/kg IV), or paclitaxel (15 mg/kg IV) + ISRIB (2.5 mg/kg IP). **i)** PDX401 tumor volumes in mice treated every second day for 2 weeks with DMSO, ISRIB (2.5 mg/kg IP), paclitaxel (20 mg/kg IV), or paclitaxel (15 mg/kg IV) + ISRIB (2.5 mg/kg IP). Arrows indicate cessation of treatment. Data presented as mean tumor volume ± SD (n=6). **j)** The number of spheres formed from 9600 viable PDX401 cells dissociated from 1cm diameter tumors taken from mice in k) and then cultured in single cell suspensions. Bars represent the mean tumorsphere count ± SD (n=6). Images of representative spheres are presented. Micron bars = 100 μm. **k)** Kaplan-Meier plot demonstrating the correlation between PERK expression (top 50% of expressers versus the bottom 50%) and survival. High PERK expression correlates with poor overall survival by Cox regression analysis when the expression level is dichotomized by median and when used outright as a continuous variable (based on stratification around median expression HR 1.80 (1.24-2.62), p = 0.0019; as a continuous variable HR 1.001 (1-1.001), p = 0.00059). Expression of PERK (eIF2αK3) transcript increases relative to normal adjacent breast tissue. EIF2α decreases in tumors relative to normal adjacent breast tissue. The abundance of both transcripts was normalized and estimated using RSEM. The hazard ratio demonstrates a significant risk to patients with higher levels of PERK compared to their low PERK-expressing cohort. For graphs, values indicated by an asterisk (*) are statistically different from controls and values indicated by letters are statistically different from each other (p<0.05).

Considering its beneficiary effects in hindering hypoxia- or INK128-induced breast cancer plasticity, we next tested whether ISRIB may improve the efficacy of paclitaxel. We exposed T47D and SUM149 cells to varying concentrations of paclitaxel (2.5-50nM) in the presence or absence of ISRIB (10nM) and then measured surviving fractions using colony formation assays. We determined that ISRIB sensitizes both cell lines to paclitaxel (Figure 5c; Supplemental Figure 5b). We next extended these cell culture studies *in vivo* to include established SUM149 xenografts in the mammary fat pads of mice. When SUM149 tumors reached a diameter of 5mm, we treated mice with vehicle (DMSO), ISRIB (2.5mg/kg IP every second day), paclitaxel (15mg/kg IV, weekly), or paclitaxel/ISRIB combination therapy for 2 weeks. IHC on tumor sections at endpoint confirmed that ATF4 levels were increased by paclitaxel, which was reversed by ISRIB (Supplemental Figure 5c). All treatment groups exhibited a reduction in tumor growth rate, with paclitaxel markedly reducing tumor burden in the presence or absence of ISRIB (Figure 5d). Cancer cells dissociated from tumors grown in paclitaxel-treated animals had higher frequencies of stem cell-like cells than did those treated with vehicle, whereby this effect of paclitaxel was mitigated by ISRIB (Figure 5e). Due to toxicities, breast cancer patients are often treated with suboptimal doses of paclitaxel. Accordingly, we sought to determine whether ISRIB could improve the efficacy of suboptimal chemotherapy. For these studies, SUM149 breast cancer cells were implanted into the mammary fat pads of mice and when tumors reached a diameter of 5mm, we treated mice with vehicle (DMSO), ISRIB (2.5mg/kg IP), paclitaxel (20mg/kg IP), or paclitaxel/ISRIB combination therapy for 2 weeks. Using this paradigm, we found that paclitaxel reduced tumor growth to the same extent as ISRIB and that ISRIB improved the efficacy of paclitaxel (Figure 5f). Moreover, histological examination demonstrated that ISRIB reduced cell viability within the necrotic regions of tumors (Figure 5g). We confirmed these results using another triple negative breast cancer model (MDA-MB-231). Mice bearing MDA-MB-231 tumors were treated with vehicle (DMSO), paclitaxel (15mg/kg IV), or paclitaxel/ISRIB (2.5mg/kg IP) combination therapy and then tumor volumes were calculated one week after cessation of therapy. ISRIB dramatically improved the efficacy of paclitaxel in this model (Figure 5h). This is in accordance with a recent study showing that ISRIB is particularly effective at reducing tumor burden in metastatic prostate cancers, which are characterized by high levels of phospho-eIF2α ^50^.

We next set out to determine whether ISRIB could also improve the efficacy of paclitaxel in clinically relevant PDX models, wherein we treated mice with paclitaxel IV and ISRIB via gavage as has been recently described ^50^. We established PDX401 and PDX574 xenografts in the mammary fat pads of mice. We chose to use a poorly differentiated triple negative breast cancer PDX (574) that contains high levels of hypoxia (marked by CA9) that co-localizes with NODAL, as well as a well differentiated PDX (401), which contains relatively little hypoxia and lower levels of NODAL (Supplemental Figure 5d) in order to account for both high and low BCSC-containing tumors. Using flow cytometry, we confirmed that NODAL is enriched in the CA9 fraction of tumors (Supplemental Figure 5e). When tumors reached a diameter of 5mm, mice were treated with DMSO, ISRIB (10mg/kg orally daily for 2 weeks), paclitaxel (15mg/kg iv weekly for 2 weeks), or the combination therapy of ISRIB and paclitaxel. IHC on tumors at endpoint confirmed that ATF4 levels were increased by paclitaxel, whereas ISRIB prevented these effects (Supplemental Figure 5f,g). While PDX 574 mice were resistant to paclitaxel, ISRIB decreased tumor growth in these mice (Supplemental Figure 5h). Cancer cells dissociated from tumors grown in paclitaxel-treated animals had higher frequencies of cells exhibiting stem cell-like properties than did those treated with vehicle, and this effect of paclitaxel was dramatically attenuated by ISRIB (Supplemental Figure 5i). In turn, paclitaxel completely eradicated tumor growth in paclitaxel-sensitive PDX401 mice (Figure 5i). Nonetheless, cancer cells dissociated from PDX401 tumors grown in paclitaxel-treated animals still exhibited higher BCSC frequencies than did those treated with vehicle, but this effect of paclitaxel was attenuated by ISRIB (Figure 5j). Notably, these alterations in BCSC frequencies occurred even when tumor growth differences were not observed, suggesting that ISRIB may selectively prevent the acquisition of BCSC-associated phenotypes.

Collectively, using several models, we demonstrated that ISRIB can attenuate the induction of plasticity in breast cancer induced by a variety of stimuli including hypoxia, mTOR inhibitors and paclitaxel. Accordingly, analysis of RNA sequencing data from 1100 breast cancer patients in the TCGA demonstrates that high levels of PERK, which phosphorylates eIF2α during the ISR, are predictive of poor survival for 10 years following diagnosis with a hazard ratio of approximately 1.8 (Figure 5k).

## Discussion

We demonstrate that *NANOG*, *SNAIL* and *NODAL* mRNAs exist in multiple isoforms which differ in their 5’UTRs, and that the selective translational induction of a subset of these isoforms underpins the acquisition of breast cancer cell plasticity in response to hypoxia, mTOR inhibition and chemotherapy. This is achieved via specific 5’UTR features present in a subset of *NANOG*, *SNAIL* and *NODAL* mRNA isoforms which allow their efficient translation under conditions wherein translational apparatus is perturbated by cancer-related stresses including hypoxia and chemotherapeutics. Importantly, the induction of plasticity through selective translation enhances tumorigenicity by increasing CSCs, leading to increased metastasis and therapy resistance. Moreover, these types of stresses are an intrinsic aspect of the tumor microenvironment (e.g. hypoxia) that are further amplified by chemotherapy. We show that by inhibiting pluripotency factors like NODAL, or by interfering with the ISR, the plastic adaptation to stress can be circumvented. These data provide previously unrecognized critical insights into the mechanisms which underpin the effects of niche microenvironmental stresses on the acquisition of tumorigenic phenotypes and the accumulation of cancer stem cell-like properties.

It is thought that mRNAs are translationally activated or inactivated in response to environmental stimuli and/or intercellular cues based on their 5’UTR features, which are considered to be static ^14^. For example, the translation of uORF containing mRNAs such as *ATF4* is low at baseline and stimulated by stress, whereas TOP mRNAs are translationally activated by nutrients and are suppressed when nutrients are limited {Ivanov, Hinnebusch, Sonenberg}. In this model, translational activity is mediated by changes in the translational machinery, rather than by the selection of mRNA isoforms. For instance, induction of eIF2α phosphorylation and consequent reduction in ternary complex (TC) availability favors translation of uORF containing transcripts such as *ATF4* from their major uORF because translation from inhibitory uORFs is not engaged when TC levels become limiting ^14^. In turn, stimulation of cells with nutrients, growth factors and hormones activates mTOR, which in addition to increasing global protein synthesis, reprograms the translational machinery to bolster translation of mRNAs harboring TOP motifs, or 5’UTRs of non-optimal length ^51–53^. In attempting to understand the mechanisms by which stem cell factors are selectively translated, we made the surprising observation that translation of mRNAs can be regulated in cis, due to the existence of multiple transcriptional start sites (TSSs) and thus mRNA isoforms (each with different 5’UTR features). Due to differences in 5’UTR sequence, each isoform has a distinct sensitivity to alterations in the translational machinery, which leads to differential translation of specific isoforms under different stress conditions. Accordingly, translational activation is enabled under a wide array of cellular states, whereby dynamic selection of 5’UTRs rather than static 5’UTR properties, dictates translational activity. Our study illustrates that this diversity of 5’UTRs in the isoforms of the mRNAs that encode factors that drive plasticity, enables their translation under stress and that this translation drives the induction of BCSCs. It will be interesting to determine how pervasive this phenomenon is; and to establish whether transcriptional start site selection may alter the ratios of each transcript, in a way that may either counteract or synergize with alterations in the translational machinery.

MTOR inhibitors have been exploited clinically to target aberrant translation in neoplasia ^54^. To this end, rapamycin and its analogues (rapalogs) have been approved for treatment of a subset of cancer patients, including those suffering from breast cancer, whereas a new generation of active site mTOR inhibitors is undergoing numerous clinical trials ^42^. In this study, we have revealed that mTOR inhibitors unexpectedly induce breast cancer cell plasticity by inducing the translation of NANOG, SNAIL and NODAL isoforms in a manner similar to hypoxia. This appears to occur via mTOR-ISR cross-talk and is likely to contribute to the lesser than expected clinical efficacy of using mTOR inhibitors in breast cancer and other oncological indications. Indeed, much like chemotherapies, mTOR inhibitors would be expected to reduce the ability of primary tumors and metastases to grow; however, under chronic treatment an unintended consequence may be the propagation of a more aggressive disease, fueled by the accumulation of BCSCs. In illuminating the mechanism by which the mTOR-mediated enhancement of BCSCs occurs, this study also revealed that mitigating the ISR with a compound such as ISRIB can prevent the adaptive acquisition of BCSC phenotypes. Given the role of BCSC-associated plasticity in therapy resistance and metastasis, these results suggest that administration of ISRIB may help improve outcomes in breast cancer patients treated with mTOR inhibitors and/or chemotherapy.

In conclusion, we have illuminated a previously unappreciated mechanism of translational regulation whereby plasticity-inducing factors are translated during stress due to the presence of multiple mRNA isoforms harboring diverse 5’UTR sequences. We have further demonstrated that inhibiting this process, with the use of ISRIB; can prevent the acquisition of BCSCs in hypoxic tumors or in response to therapy, so that adaptive therapy resistance and/or metastatic dissemination may be prevented.

## METHODS

### Cell Culture and Treatments

T47D, MCF7 and MDA-MB-231 cells, obtained from ATCC (Manassas, Virginia, USA), were maintained in RPMI-1640 Medium (Life Technologies; Carlsbad, California, USA) with 10% fetal bovine serum (FBS) (Life Technologies). Cells were passaged using 0.25% (w/v) Trypsin (Life Technologies) as per ATCC recommendations. SUM149 cells, purchased from Bioreclamation IVT, were grown in Ham’s F-12 medium with 5% heat-inactivated FBS, 10mM HEPES, 1µg/mL hydrocortisone, 5µg/mL insulin. Breast cancer cells were authenticated at the Sick Kids Research Institute and tested for mycoplasma in house. H9 hESCs from WiCell (Madison, Wisconsin, USA) grown on irradiated CF-1 Mouse Embryonic Fibroblasts (GlobalStem; Gaithersburg, Maryland, USA) in knockout DMEM/F12 (Life Technologies; Carlsbad, California, USA), 20% knockout serum replacement (Life Technologies), 1X non-essential amino acids (Life Technologies), 2mM glutamine (Life Technologies), 0.1mM 2-mercaptoethanol (BME; Thermo Fisher Scientific; Waltham, Massachusetts, USA), and 4ng/mL of basic fibroblast growth factor (FGF) (Life Technologies). For experiments, cells were passaged into feeder-free conditions. Feeder-free conditions consisted of Geltrex matrix (Life Technologies) as a growth substrate and mTeSR1 media (Stem Cell Technologies; Vancouver, British Columbia, Canada). All cells were grown in a humidified environment at 37°C with 5% CO_2._

#### Hypoxia

Hypoxia was administered at the noted concentrations using the Xvivo system (BioSpherix; Parish, New York, USA). Temperature (37°C) and CO_2_ (5%) were maintained.

#### Manipulation of NODAL

To increase Nodal signaling, we used a Nodal expression vector (versus an empty pcDNA3.3 vector; pcDNA™3.3-TOPO® cloning kit; Invitrogen) as previously described^23, 55^. We also employed recombinant human Nodal (rhNodal; R&D). To decrease Nodal signaling, we used Nodal-targeted shRNAs (versus scrambled control shRNAs) as previously described^23, 55^. Transfection was performed with Lipofectamine (Invitrogen) as per manufacturer instructions. For stable selection, Puromycin (200-450 ng/mL) or Geneticin (G418; 800 ng/mL) was used. To inhibit Nodal signaling, we also used SB431542. SB431542 selectively inhibits Activin and TGF-β and Nodal signaling but not BMP signaling.

#### Manipulation of mTOR and 4EBP1

To inhibit mTOR kinase activity we used MLN0128 (INK; 20nM). To increase 4E-BP1 levels we used an expression vector (pCMV6-Entry EIF4EBP1 True ORF Gold Vector; OriGene). To knock down 4E-BP1 we used pGFP-V-RS EIF4EBP1 Human shRNA (OriGene) versus scramble and control vector. Transfection was performed with Lipofectamine 2000 (Invitrogen) as per manufacturer instructions.

#### Manipulation of ISR components

To induce ER stress, Azetidine (AZE; 5mM) or Thapsigargin (TG, 0.1 μM) were used. Paclitaxel (Pac; 20nM) was also used. In order to examine the role of eIF2α phosphorylation in the hypoxic induction of BCSC phenotypes, control (EV) MDA-MB-231 cells or cells wherein eIF2α was knocked down and the replaced with either a wild type (WT) eIF2α or a mutant (KI) eIF2α that cannot be phosphorylated were used as previously described^46^. In order to prevent eIF2α dephosphorylation, Salubrinal (20μM) was used. In order to overcome the effect of eIF2a on ternary complex turnover and translation, the Integrated Stress Response Inhibitor (ISRIB; 10 nM) was used.

### Western Blotting

Cells were lysed on-plate using Mammalian Protein Extraction Reagent (M-PER; Thermo Scientific), with Halt Protease Inhibitor Cocktail (Thermo Scientific), and Phosphatase Inhibitor (Thermo Scientific). Protein was quantified according to manufacturer’s instructions utilizing Pierce BCA Protein Assay Kit (Thermo Fisher) and measured on a FLUOstar Omega plate reader (BMG LABTECH; Offenburg, Germany). 4x Laemmli buffer (Bio-Rad; Hercules, California, USA) with 5% BME (Sigma-Aldrich; St. Louis, Missouri, USA) containing 20µg of protein was boiled for 10 minutes and loaded to be analyzed. Samples were separated by SDS-polyacrylamide gel electrophoresis, and then transferred onto Immobilon-FL membranes (Millipore). Precision Plus Protein Dual Color Standards (Bio-rad) was used to approximate molecular weight. Membranes were blocked with 5% milk in PBS 0.1% Tween (Sigma-Aldrich) for 1 hour at room temperature, then incubated with primary antibody overnight at 4°C (Supplementary Table 2). After washing in PBS 0.1% Tween (Sigma-Aldrich), membranes were incubated with horseradish peroxidase-conjugated secondary antibodies (Bio-Rad) and then washed to remove excess secondary antibody. Clarity Western ECL Substrate (Bio-Rad) was used to detect signal. ChemiDoc™ XRS+ System (Bio-Rad) or film were used to image the western blots. Densitometry was performed using ChemiDoc™ XRS+ System (Bio-Rad).

#### Florescence western blot detection

Using the Trans Blot Turbo (settings of 25 V and 1.3 A for 15 minutes; Bio-rad) proteins were transferred to a low-auto-fluorescence PVDF membrane (Bio-rad), blocked for one hour at room temperature with Odyssey Blocking Buffer (Li-Cor; Lincoln, Nebraska, USA), then incubated with primary antibody overnight at 4°C in Odyssey Blocking Buffer with 0.1% Tween-20 (Sigma-Aldrich). Membranes were then probed with corresponding Li-Cor anti-mouse or anti-rabbit fluorescent secondary antibodies for one hour at room temperature at dilutions of 1/10 000 in Odyssey Blocking Buffer with 0.1% Tween-20 (Sigma-Aldrich) and 0.1% Tween. Imaging was conducted using the Li-Cor Odyssey Clx imaging system. Scans were performed at intensities that did not result in any saturated pixels.

### Polysome Profiling

Cells were grown to 60-80% confluency at which time 0.1 mg/ml of cycloheximide was added to cells for 5 min at 37 °C before harvesting. The cells were extracted in polysome lysis buffer (15 mM Tris·HCl (pH 7.4)/15 mM MgCl2/0.3 M NaCl/1% Triton X-100/0.1 mg/ml cycloheximide/100 units/ml RNasein), and the volume of each lysate to be loaded onto gradients was determined by total RNA. Sucrose gradients (7-47%) were centrifuged at 39,000 rpm with a SW41-Ti Rotor (Beckman Coulter, Fullerton, CA) for 90 min at 4 °C. Gradients were continuously monitored at an absorbance of 254 nm and fractionated with a Brandel BR-188 Density Gradient Fractionation System. Each gradient was collected into nine equal fractions. The baseline absorbance of the sucrose gradient was calculated from the absorbance of a blank gradient using Peakchart software and subtracted from the absorbance reading of each sample. RNA isolation was conducted by first digesting each fraction with proteinase K, and extracting total RNA by phenol-chloroform extraction and ethanol precipitation. Samples were pooled into groups representing monosomes, light (less than 3 ribosomes) and heavy (more than 3 ribosomes) polysomes. Equal amounts of RNA were analyzed by real-time RT PCR. For validation studies, mRNA in polysomes (more than 3 ribosomes) was compared to total mRNA levels. For polysome profiles, the percentage of transcript in each fraction was calculated.

#### RNA Extraction and RT-PCR

The PerfectPure RNA Cultured Cell Kit (5-Prime; Hilden, Germany) was used to extract total RNA from cultured cells following the manufacturer’s protocol. Optional DNase treatment was performed, and RNA was eluted in 50µL. 3µL of purified RNA was used for quantification using the Epoch plate reader (Biotek; Winooski, Vermont, USA). cDNA was made from purified total RNA using high capacity cDNA reverse transcription kit (Applied Biosystems; Foster City, California, USA) as per manufacturer’s protocol. The included random hexamers were used to prime reverse transcription and 1µg of RNA was used for each. ‘No Template’ reactions did not contain RNA and ‘No RT’ reactions did not contain reverse transcriptase enzyme. Real time PCR analysis was performed on 1µL of cDNA using TaqMan Gene Expression Master Mix according to the manufacture’s procedures using FAM labeled TaqMan® and PrimePCR gene expression human primer/probe sets (Thermo Scientific and BioRab; see Supplementary Table 3). mRNA expression was compared to untreated control using the ΔCT method. Data was collected on a CFX96 Touch Real-Time PCR Detection System (Bio-Rad; Hercules, California, USA) using standard real time PCR settings. Activation 95°C 10 min; Melting 95°C 15 sec; Annealing/ extension 55°C 1 min. Return to step 2 for 40 total cycles. Melt curve analysis was performed to ensure the production of a single amplicon. For absolute quantification of NODAL, after cDNA synthesis real time PCR was performed using Power SYBR Master Mix (Life Technologies). 1µL of the cDNA was loaded in triplicate for quantification of NODAL (1µL = 50ng of starting RNA). The following primers were used: NODAL forward primer TACATCCAGAGTCTGCTG; and NODAL reverse primer CCTTACTGGATTAGATGGTT. Cloned Nodal PCR products were linearized, and diluted series was made (copy number/µL). A standard curve was constructed from these samples and run with the samples to estimate the number of NODAL transcripts at each time point. For isoform specific PCR all products were validated using sequencing. The following primers were used: NANOG 350 forward primer GAT GGG GGA ATT CAG CTC AGG; NANOG 350 reverse primer TCA AGA CTA CTC CGT GCC CA; NANOG 291 forward primer AAC GTT CTG GAC TGA GC; NANOG 291 reverse primer AGG CAG CTT TAA GAC TTT TCT GG; SNAIL 417 forward primer AAA GGG GCG TGG CAG ATA AG; SNAIL 417 reverse primer CGC CAA CTC CCT TAA GTA CTC C; SNAIL 85 forward primer CGG CCT AGC GAG TGG TTC; SNAIL 85 reverse primer CAC TGG GGT CGC CGA TTC; NODAL small (42 + 14 + 298) forward primer CTG GAG GTG CTG CTT TCA GG; NODAL small (42 + 14 + 298) reverse primer CAG GCG TGC AGA AGG AAG G; NODAL 298 forward primer GTT TGG TAC CTA GAG CAG G; NODAL 298 reverse primer TCC AGG GAC GGG ATC TAG G; NODAL 416 forward primer CCC TCG GCA TTC TCT TCC TG; NODAL 416 reverse primer ATC CCT GCC CCA TCC TCT C.

### Sphere Formation

Sphere formation media was composed of DMEM/F12 + GlutaMax (Life Technologies), 1x B27 (Life Technologies), 20 ng/mL epidermal growth factor (EGF) (Life Technologies), and 10 ng/mL FGF (Life Technologies). After treatment, cells were harvested using 0.25% (w/v) trypsin (Life Technologies), the trypsin was neutralized, and the cells resuspended in fresh media. These cells were filtered through a 40µm pore filter (Thermo Fisher) to obtain a single cell solution. Cells were counted using trypan blue and diluted in the sphere formation media to the appropriate concentration for plating. 200µL of the diluted cells were seeded into each well of a 96 well Ultra-Low Attachment Surface plate (Corning, NY, USA). Spheres were given between 10 and 21 days to grow. Images of spheres were captured using the EVOS FL Cell Imaging System (Thermo Fisher) at 4X magnification. In order to enrich for spheres, cells were cultured in a bioreactor (Synthecon) as previously described^56^.

### Flow Cytometry

One million cells were stained in 100µL of Zombie Aqua (Fixable Viability Kit BioLegend; San Diego, California, USA) for twenty minutes at room temperature. Zombie aqua was removed and 20µL of antibody dilution was added to each sample, which was then incubated on ice for 10-15 minutes. NODAL and CA9 staining was performed on fixed and permeabilized cells according to manufacturer instructions using the Fixation/Permeabilization kit (BD Biosciences).

Antibody Pairs:

– CD24 APC (REA832, MiltenyiBiotec, 1:20 dilution), CD44 Vioblue (REA690, Miltenyi Biotec, 1:5 dilution) used in Figure 1f
– FITC Mouse Anti-Human CD24 (BD Biosciences; Franklin Lakes, New Jersey, United States, 1:5 dilution), PE Mouse Anti-Human CD44 (BD Biosciences, 1:5 dilution) used in Figure 3d and Supplemental Figures 1b-d and 3f,g
– Human Carbonic Anhydrase IX/CA9 Fluorescein-conjugated Antibody (R&D Systems, Minneapolis, Minnesota, United States, 1:20 dilution)
– Human NODAL AF647 (Novus Biologicals, Basel, Switzerland, 1:20 dilution)

Cells were washed with 200µL FACs buffer (PBS with 1% FBS), pelleted and then resuspended in 100ul 2% PFA in FACs buffer. For acquisition, cells were re-suspended in 300µl FACS buffer for flow acquisition. Doublet discrimination and live cell gates were used to identify the cells of interest, and quadrant gates were set according to the fluorescence minus one controls (FMO).

### RNA-seq and Gene Set Enrichment

RNA was extracted from hypoxia treated cells with the Qiagen RNeasy kit and quantified via Nanodrop; quality was measured using Qubit. RNA was shipped to McGill University and Genome Quebec Innovation Center, where quality was validated via Bioanalyser, followed by NEB/KAPA library preparation and sequencing via Illumina HiSeq. Post sequencing quality check of reads was performed with FastQC and adapter sequences removed using Skewer. Reads were aligned to the GRCh38/Hg19 human reference genome using STAR. Data processing included Bigwig, PCA, correlation matrices, and coverage maps of aligned reads were produced using DeepTools. Read quantification via FeatureCounts was performed using Refseq annotations. Expression values for paired samples, hypoxia/normoxia, were obtained using the exact test within the edgeR package. An adjusted p-value (FDR) of 0.05 was used to determine statistically significant differences (p-value adjusted for multiple hypothesis testing by the Benjamini-Hochberg method). GAGE package for R was used to compare data to the hallmark gene sets from Molecular Signatures Database, which was used for Gene set analysis.

### Luciferase Reporter Assays

For cloning of the luciferase 5’ UTR reporters, we modified the PTK-ATF4-Luc plasmid from (Dey et al., 2010)^39^ using the QuikChange Lightning Site-Directed Mutagenesis Kit (Agilent; Santa Clara, California, USA) to introduce two BsmBI restriction sites flanking the ATF4 5’ UTR. The ATF4 5’ UTR was then replaced with a LacZ insert to complete the pGL3-TK-5UTR-BsmBI-Luciferase plasmid used for one-step cloning of various 5’ UTRs. This plasmid has been made available on Addgene (plasmid #114670). Except for “Snail 417”, 5’ UTRs of interest flanked by adapter sequences for cloning were ordered as gBlocks from Integrated DNA Technologies (Coralville, Iowa, USA; sequences are listed in Supplementary Table 4) and cloned into pGL3-TK-5UTR-BsmBI-Luciferase using BsmBI (New England BioLabs; Whitby, Ontario, Canada). The “Snail 417” UTR insert was generated by PCR using the forward primer TATCGTCTCAACACCGAGCGACCCTGCATAAGCTTGGCGCTGAGCCGGTGGGCG and the reverse primer ATACGTCTCTCTTCCATAGTGGTCGAGGCACTGGGGTCG. The “NODAL 298” uORF mutant was generated using the QuikChange Lightning Site-Directed Mutagenesis Kit with the forward primer CCTCCGGAGGGGGGTTATATAATCTTAAAGCTTCCCCAG and the reverse primer CTGGGGAAGCTTTAAGATTATATAACCCCCCTCCGGAGG, to introduce a G-to-A mutation within an upstream start codon (ATG) at position −104 relative to the translational start. HEK293 cells were transfected using Polyethylenimine (PEI,Sigma Aldrich; St. Louis, Missouri, USA). PEI and vector media were combined in a ratio of 5µL PEI and 0.5µg vector in 250µL of serum-free DMEM and incubated for 10 min at room temperature. 250µL of the mixture was apportioned into each well of a 12-well plate containing 250µL DMEM. HEK293 cells were incubated overnight in the transfection mixture. The transfection media was removed and replaced with DMEM 10% serum and the cells were given 24 hours to recover. Cells were then treated with 0.1 µM of thapsigargin (TG) for 3 hours or 0.5% O_2_ for 6 hours (hypoxia). Upon completion of treatment, cells were lysed and the luciferase activity measured with the Firefly Luciferase Assay System (Promega, Madison, Wisconsin, USA). Activity was read using a FLUOstar Omega plate reader (BMG LABTECH).

### Animal Models

All experiments involving animals were approved by the Animal Use Subcommittee at the University of Alberta (AUP00001288 and AUP00001685).

#### Experimental Metastasis Assay

SUM 149 cells were pre-incubated as described, then trypsinized and counted. 500,000 cells in 700 uL Ca2+-free HBSS were injected into the tail vein of female NOD-scid IL2Rgamma^null^ (NSG) mice. Mice were sacrificed at 8 weeks (to tumor formation). Lungs were formalin-fixed and paraffin-embedded, and immunohistochemical staining on this tissue was conducted using a human-specific HLA antibody (Supplementary Table 1) as per the manufacturer’s instructions. For each mouse organ, 3-6 sections were acquired from evenly spaced areas throughout the tissue, and the average number of metastases per mouse organ was calculated.

#### Orthotopic Xenografts

500,000 SUM 149 or MDA-MB-231 cells in 100 µL RPMI:Matrigel (1:1) were injected into the right thoracic mammary fat pad of 7-8 week old female NSG mice. Mice were randomized and treatments were administered when tumors reached a maximum diameter of 5 mm. At this point, mice were treated with DMSO vehicle control, INK (30 mg/kg by gavage), ISRIB (2.5 mg/kg IP) or paclitaxel (20 mg/kg IP or 15 mg/kg IV) for the times indicated. Tumor measurements were taken twice per week and a digital caliper was used to measure Length x Width x Depth of the tumor upon excision in order to calculate volume. Mice were sacrificed when tumors reached ∼1 cm in diameter. Tumors were cut in half. One half was dissociated and the other was fixed with 4% formaldehyde, paraffin embedded, sectioned and stained with H&E or used for immunohistochemistry. Survival curves for overall survival were constructed using the Kaplan-Meier method and significance determined by log-rank test.

#### Patient Derived Xenografts

Two PDX models obtained through a collaboration with Oncotest (Charles River, Freiburg, Germany) were used: PDX 401 is a well differentiated basal-like TNBC and PDX 574 is a poorly differentiated basal-like TNBC. Viable pieces (∼1mm in diameter) were placed, through a small incision, into the mammary fat pads of 7-8 week old female NSG mice. At this point, mice were treated with DMSO vehicle control, INK (30 mg/kg by gavage), ISRIB (2.5 mg/kg IP or 10 mg/kg by gavage) or paclitaxel (20 mg/kg IP or 15 mg/kg IV) for the times indicated. Tumor measurements were taken twice per week and a digital caliper was used to measure Length x Width x Depth of the tumor upon excision in order to calculate volume. Mice were sacrificed when tumors reached ∼1 cm in diameter. One half was dissociated and the other was fixed with 4% formaldehyde, paraffin embedded, sectioned and stained with H&E or used for immunohistochemistry. Survival curves for overall survival were constructed using the Kaplan-Meier method and significance determined by log-rank test.

#### Tumor Dissociation

Half of each tumor was dissociated using the Human Tumor Dissociation Kit and Gentle MACS Tissue Dissociator with Heaters (Miltenyi Biotec; Bergisch Gladbach, Germany) according to the manufacturer’s instructions prior to enumeration of live cells using trypan blue.

#### Analysis of tumor necrosis

Three tumor sections spaced evenly throughout each tissue block were stained with H&E. Each tumor section was imaged such that the entire section was visible in one field of view. ImageJ software was used to outline and quantify the total tumor area, and the area of necrosis. Necrosis was calculated as a percentage of the total tumor area.

#### Immunohistochemistry

Formalin-fixed, paraffin-embedded tissue underwent deparaffinization in xylenes, hydration through an ethanol series, antigen retrieval with citrate buffer, and peroxidase and serum-free protein blocking. Nodal, CA9, p4E-BP, 4E-BP or ATF4 specific antibodies (Supplementary Table 2) were applied. Slides were rinsed in TBS-T, and treated with Envison+ HRP anti-mouse IgG (Dako). Color was produced with DAB (brown) substrate and counterstained with Mayer’s haematoxylin. Samples were dehydrated in reagent grade alcohol and cover slipped with permanent mounting medium. Negative control reactions were conducted with mouse IgG, isotype controls used at the same concentration as the primary antibodies.

### Analysis of Patient Data

#### Datasets

Level 3 TCGA RNAseqV2 BRCA gene expression data and clinical information was obtained from the TCGA Data Portal in August 2014. RNA-sequencing RSEM values were used in downstream analyses.

#### Data Preparation

For TCGA RNA-seq samples, relative abundance (transcripts per million, TPM) was calculated by multiplying the scaled estimate data by 106 and used in downstream analysis.

#### Statistical analysis

We conducted all analyses and visualizations in the RStudio programming environment (v0.98.501). R/Bioconductor packages ggplot2, plyr, pROC, survival, GAGE and limma were used where appropriate. 4E-BP1 and PERK expression was dichotimized with receiver operating characteristics (ROC) curves to determine the optimal cutoff for the endpoint of overall survival censorship. Quantitative differences between high versus low expression cohorts were evaluated with a student’s t-test; qualitative differences were evaluated using a Fisher exact test. Survival curves for overall survival were constructed using the Kaplan-Meier method and significance determined by log-rank test/Wilcoxon test.

### Surviving Fraction Assays

Cells were incubated with paclitaxel +/- ISRIB for 1 hour in a standard CO_2_ incubator and then washed with phosphate buffered saline (PBS) and made into a single cell suspension and plated. After 7 to 14 days, colonies were fixed with acetic acid-methanol (1:4) and stained with crystal violet (1:30).

### Statistics

Analysis was conducted using GraphPad Prism 7. Student’s *t*-test was employed for direct analysis of a single condition to the appropriate control with paired two-sample t-tests used where appropriate. P values less than 0.05 were considered statistically significant (signified by *). To analyse the relationship between multiple conditions, one-way ANOVAs for all pairwise comparisons with the Bonferroni and Holm post-hoc test were employed to detect statistically significant differences between groups and correct for family-wise error rate (signified by letters). P values less than 0.05 were considered statistically significant.

## Acknowledgements

This work was supported by the Canadian Institutes for Health Research (PLS 9538 and PLS 95381), the Canadian Breast Cancer Foundation Prairies, the Alberta Cancer Foundation, the Women and Children’s Health Research Institute and Alberta Innovates Health Solutions through grants awarded to L.M.P. S.D.F. has been supported by a doctoral scholarship from the Natural Sciences and Engineering Research Council of Canada, and an Ontario Graduate Scholarship. KMV was a Vanier scholar and LL and MC were supported by PhD scholarships from the AIHS and CIHR. L.M.P. was the recipient of the premier new investigator award from the CIHR, the Sawin-Baldwin Chair in Ovarian Cancer, the Dr. Anthony Noujaim Legacy Oncology Chair and the AIHS translational health chair in cancer. IT is a scholar of the Fonds de Recherche du Québec-Santé (FRQS; Junior 2). This work was also supported by R-37-DK060596 and R01-DK 053307 awarded to MH and by CIHR MOP 38160 and a grant from the Quebec Breast Cancer Foundation awarded to AK.

## Author contributions

LMP and MJ conceived the project, wrote the manuscript and produced the figures. MJ conducted Western blots, RT-PCR assays, reporter assays, sphere formation assays, flow cytometry and transfections. LL performed quantification of mRNA on polysomes, RNAseq analysis quantification of necrosis. GZ and JL performed all animal experiments and IHC. SF designed isoform-specific PCR assays and derived constructs for reporter assays; KV conducted analyses of patient data. KT performed and validated polysome profiles, DDC measured NODAL in cells grown in 3D. DQ conducted sphere formation assays related to NODAL. ID assisted with flow cytometry. MC and ZX assisted with Western blotting and RT-PCR of stem cell genes. BJG, MH, AB and JU assisted with polysome fractionation and CP and AK provided the eIF2α KI expressing cells. JS supplied PDX models, GMS assisted with editing, flow cytometry and characterization of PDX models. IT assisted with project design and editing.

---

**Supplemental Figure 1:**
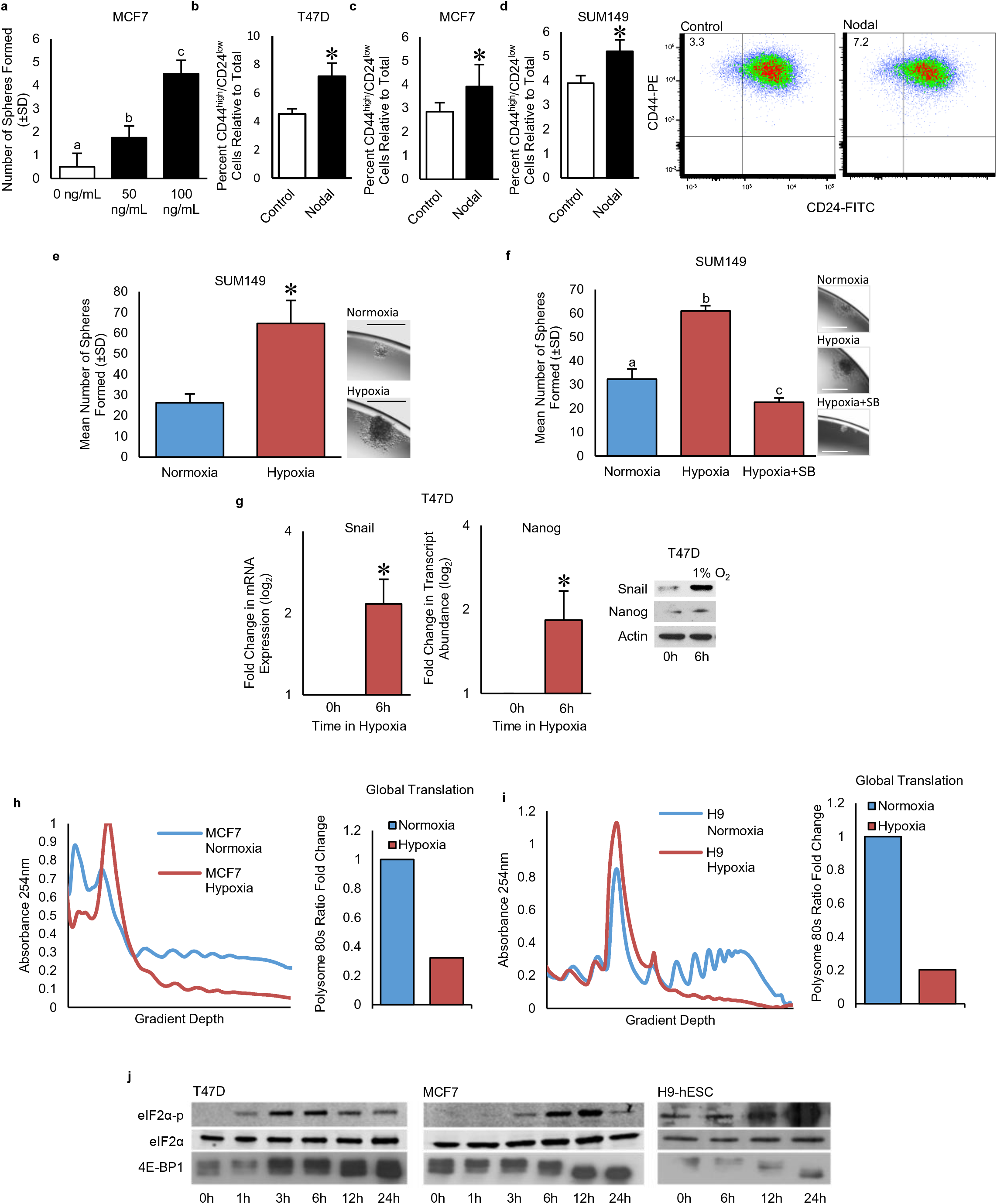
Hypoxia induces BCSC phenotypes concomitant with translational reprogramming: **a)** The number of spheres formed from MCF7 cells treated with rhNODAL (10, 100 ng/mL). Bars represent the mean tumoursphere count relative to untreated cells ± SD (n=6). **b-d) b)** T47D **c)** MCF7 and **d)** SUM149 cells expressing a CD44high/CD24low signature following exposure to rhNODAL for 24h. Bars represent mean percentage of CD44high/CD24low cells ± SD (n=6). Representative scatter plots discriminating subpopulations as defined by cell surface markers CD44 and CD24 are shown. **e)** The number of spheres formed from viable SUM149 cells pre-exposed to hypoxia or normoxia for 24h and then cultured as single cell suspensions. Bars represent the mean tumoursphere count ± SD (n=3). Images of representative spheres are presented. Micron bars = 100 μm. **f)** The number of spheres formed from viable SUM149 cells pre-exposed to hypoxia or normoxia for 24h +/- SB431542 (10 μM) and then cultured as single cell suspensions. Bars represent the mean tumoursphere count ± SD (n=3). Images of representative spheres are presented. **g)** NANOG and SNAIL transcript (detected with real time RT-PCR) and protein levels (detected with Western blotting) in T47D cells cultured in hypoxia for 0 and 6h (n=3). Bars represent Log2 fold change in transcript relative to levels at 0h and Actin is used as loading control for Western blottingTranslation rates in **h)** MCF7 and **i)** H9 cells cultured in hypoxia or normoxia for 24h. Bars represent fold change of mRNA associated with polysomes (more than 3 ribosomes) in cells cultured in hypoxia versus normoxia for representative polysome profiles shown. For graphs, values indicated by an asterisk (*) are statistically different from controls and values indicated by letters are statistically different from each other (p<0.05). **j)** Western blots of lysates from T47D breast cancer cells, MCF7 breast cancer cells and H9 hESCs exposed to 20 (normoxia) or 1% O_2_ (hypoxia) for 0-24h show that low O_2_ reduces 4E-BP phosphorylation (indicated by a downward shift) by 12h, and increases eIF2α phosphorylation between 3 and 12h. Total 4E-BP, rpS6 and eIF2α levels were unchanged.

**Supplemental Figure 2:**
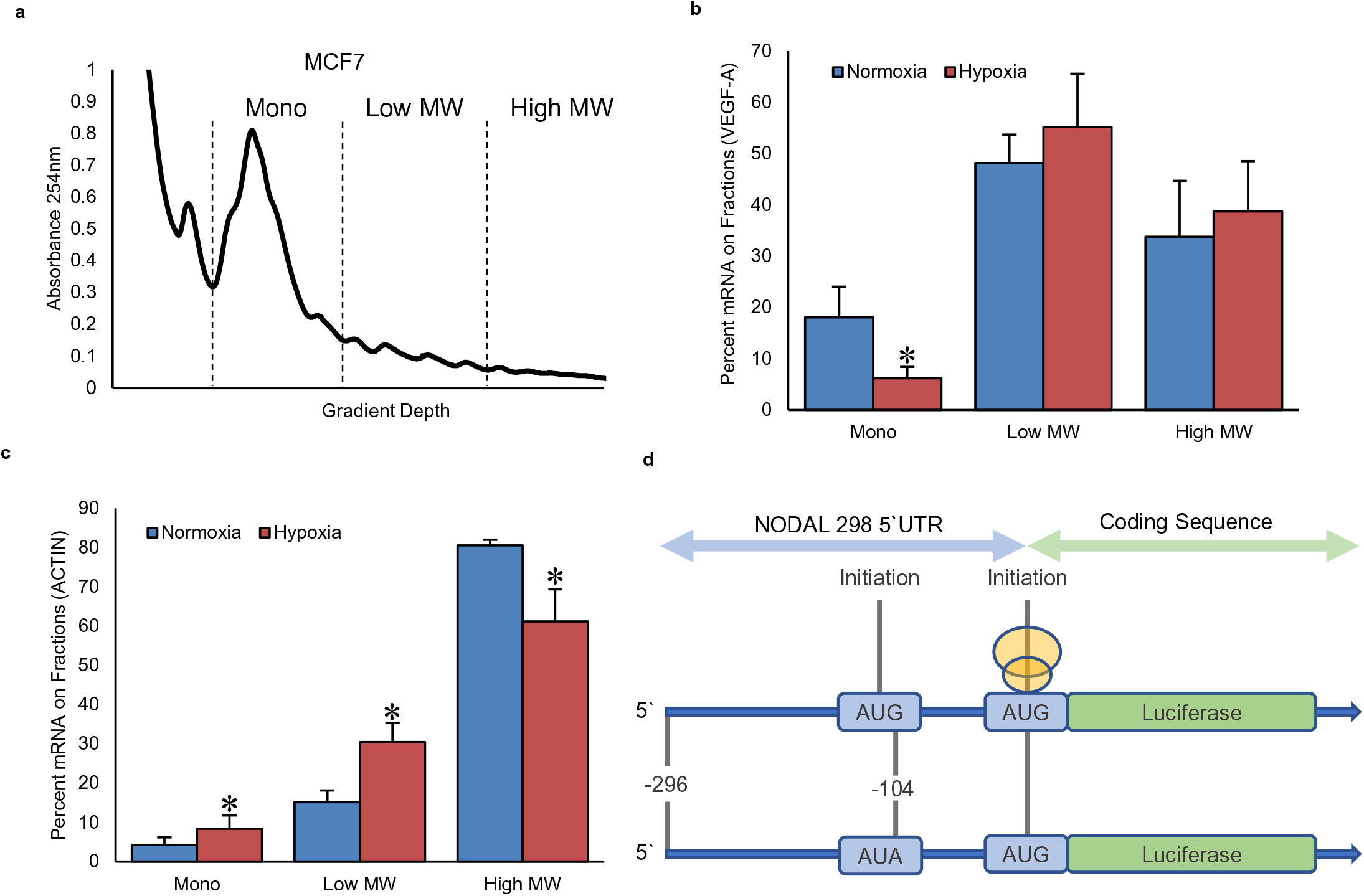
Hypoxia induces the selective translation of BCSC-associated transcripts in an isoform-specific manner. **a)** Representative polysome profile from MCF7 demonstrating the fractions from which monosomes, Low MW polysomes and High MW polysomes were taken. **b)** *ACTIN* and **c)** *VEGF* mRNA associated with mononsomes, low MW polysomes and high MW polysomes extracted from MCF7 cells cultured for 24h in hypoxia or normoxia. Bars represent the percent of transcript associated with each fraction in each condition ± SD (n=3). Values indicated by an asterisk (*) are significantly different. **d)** Sequence of the NODAL 298 nt 5’UTR highlighting the uORF.

**Supplemental Figure 3:**
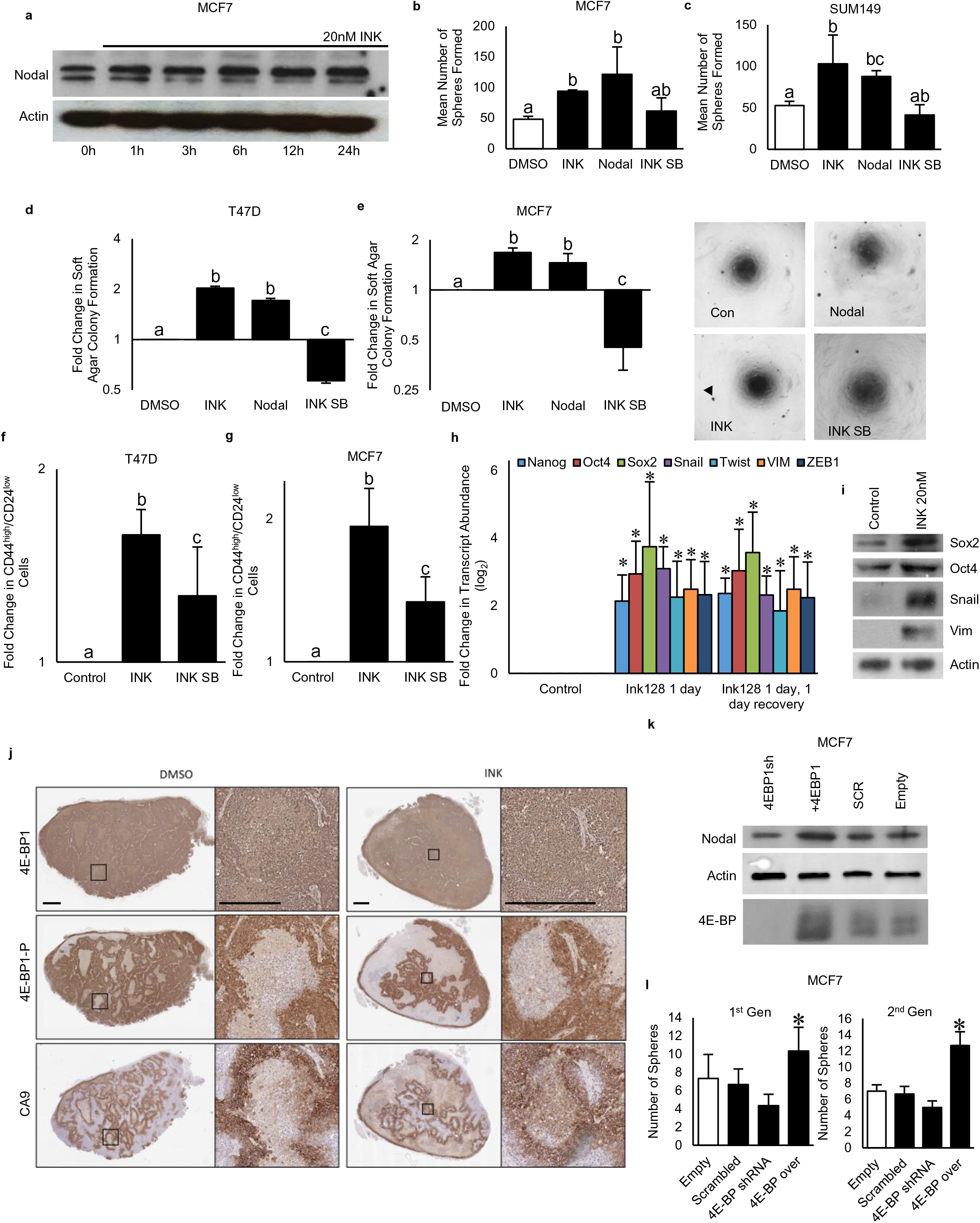
MTOR inhibition induces breast cancer plasticity: **a)** Western blot of lysates from MCF7 cells treated for 0-24h with MLN0128/INK128 (INK; 20nM). NODAL protein increases over time and Actin is used as a loading control. **b,c)** The number of spheres formed from viable **b)** MCF7 and **c)** SUM149 cells pre-exposed to DMSO, INK (20nM), rhNODAL (100 ng/mL) or INK + SB431542 (10 μM) for 24h and then cultured as single cell suspensions. Bars represent the mean tumoursphere count ± SD (n=4). Images of representative spheres are presented. **d,e)** The number of colonies formed from **d)** MCF7 and **e)** SUM149 cells pre-exposed to DMSO, INK (20nM), rhNODAL (100 ng/mL) or INK + SB431542 (10 μM) for 24h and then cultured in soft agar. Bars represent the mean colony count ± SD relative to colonies formed from cells treated with DMSO (n=3). **f,g)** Percentage of **f)** MCF7 and **g)** SUM149 cells expressing CD44high/CD24low following exposure to DMSO, INK or INK+SB431542. Bars represent mean percentage of CD44high/CD24low cells ± SD. **h)** Real time RT-PCR based quantification of transcripts associated with stem cells (NANOG, SOX2, OCT4) or EMT (TWIST, ZEB1, SNAIL, VIM) in SUM149 cells cultured for 24h in DMSO or INK (20 nM) (n=3). A group wherein INK was washed out for 24h was also included. Bars represent mean Log2 fold change relative to cells treated with DMSO ± SD. **i)** Western blot analyses of proteins associated with stem cells (SOX2, OCT4) or EMT (SNAIL, VIM) in SUM149 cells cultured for 24h in DMSO or INK (20 nM). Actin is used as a loading control. **k)** Immunohistochemical detection (brown) of phospo-4E-BP1, total 4E-BP1 and CA9 in PDX 401 tumours taken from mice treated every second day for 2 weeks with either DMSO or INK (30 mg/kg). Micron bars = 100 μm. **l)** Western blot analyses of NODAL and 4E-BP1 in MCF7 cells stably transfected with 4E-BP1 shRNA, a shRNA scrambled (SCR) control, a 4E-BP1 ORF or an empty vector control. Actin is used as a loading control. **l)** The number of spheres formed from MCF7 cells transfected with vectors as described in **k)** and then cultured as single cell suspensions. Self-renewal was further assessed by measuring sphere formation in the 2^nd^ generation. Bars represent the mean tumoursphere count ± SD. Images of representative spheres are presented. Micron bars = 100 μm. For graphs, values indicated by an asterisk (*) are statistically different from controls and values indicated by letters are statistically different from each other (p<0.05).

**Supplemental Figure 4:**
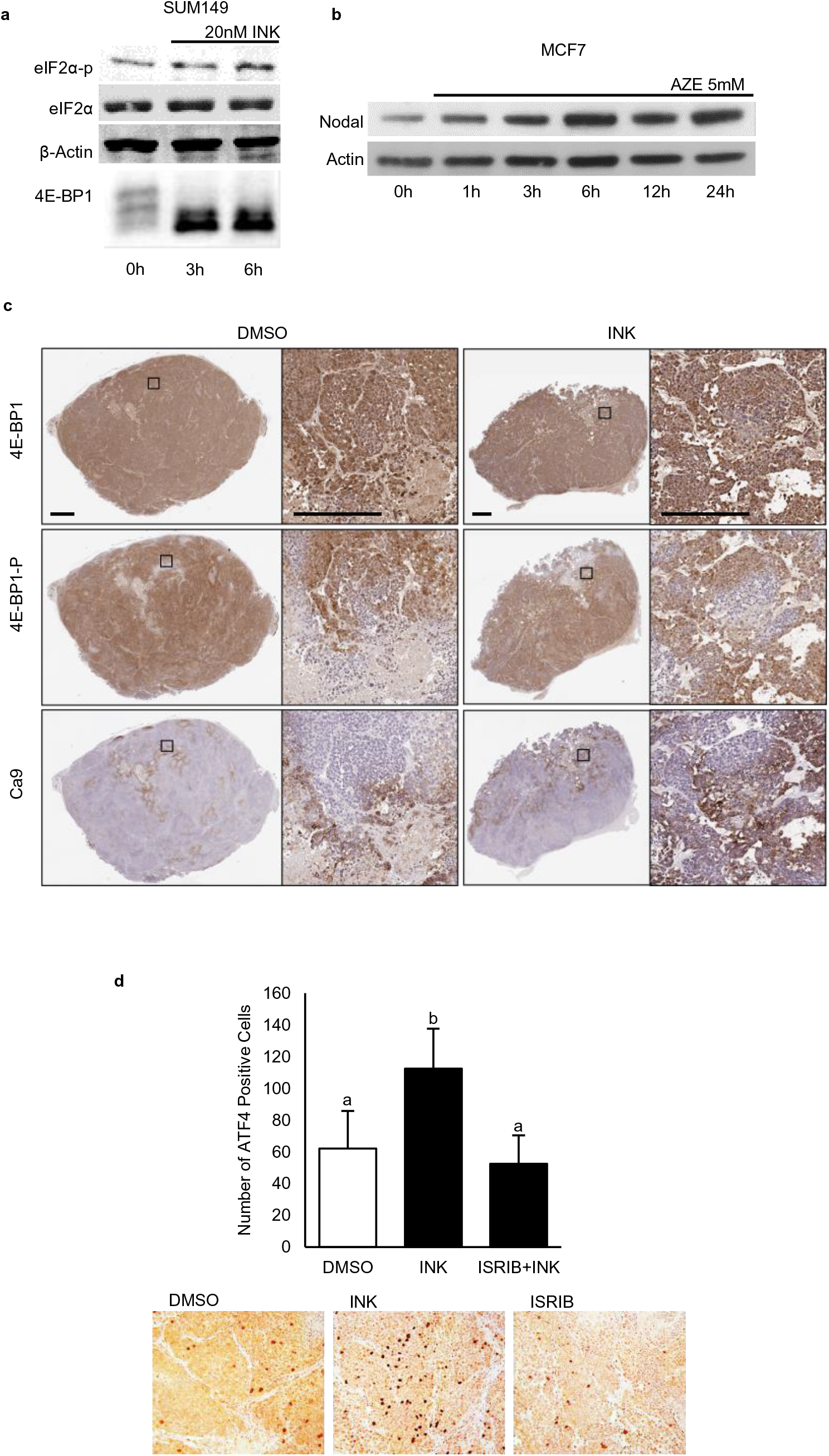
Stress-induced plasticity is mediated by the ISR: **a)** Western blots of eIF2α-p, eIF2α and 4E-BP1 in lysates extracted from SUM149 cells treated for 0, 3 or 6h with INK 20 nM show that INK induces eIF2α phosphorylation. Actin is used as a loading control. **b)** Western blot of NODAL in lysates from MCF7 cells treated with AZ (5mM) for 0-24h shows that this ER stress induces NODAL. Actin is used as a loading control. **c)** Immunohistochemical detection (brown) of phospo-4E-BP1, total 4E-BP1 and CA9 in SUM149 tumours from mice treated every second day for 2 weeks with DMSO, INK (30 mg/kg) or or INK+ISRIB (2.5 mg/kg). Micron bars = 100 μm. **d)** Immunohistochemical detection (brown) of ATF4 in SUM149 tumours from mice treated every second day for 2 weeks with DMSO, INK (30 mg/kg) or or INK+ISRIB (2.5 mg/kg) (n=3). Values indicated by letters are statistically different from each other (p<0.05).

**Supplemental Figure 5:**
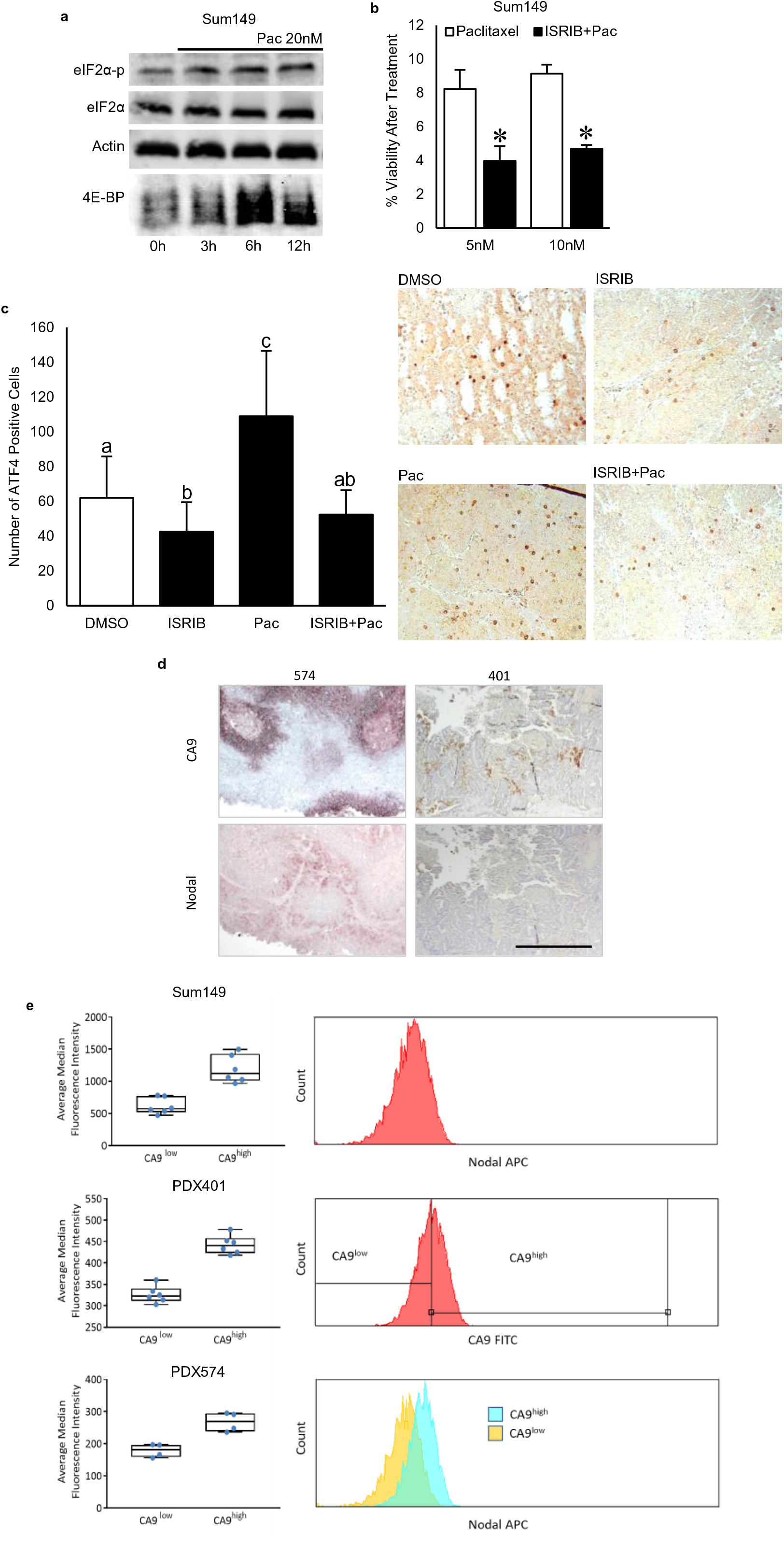
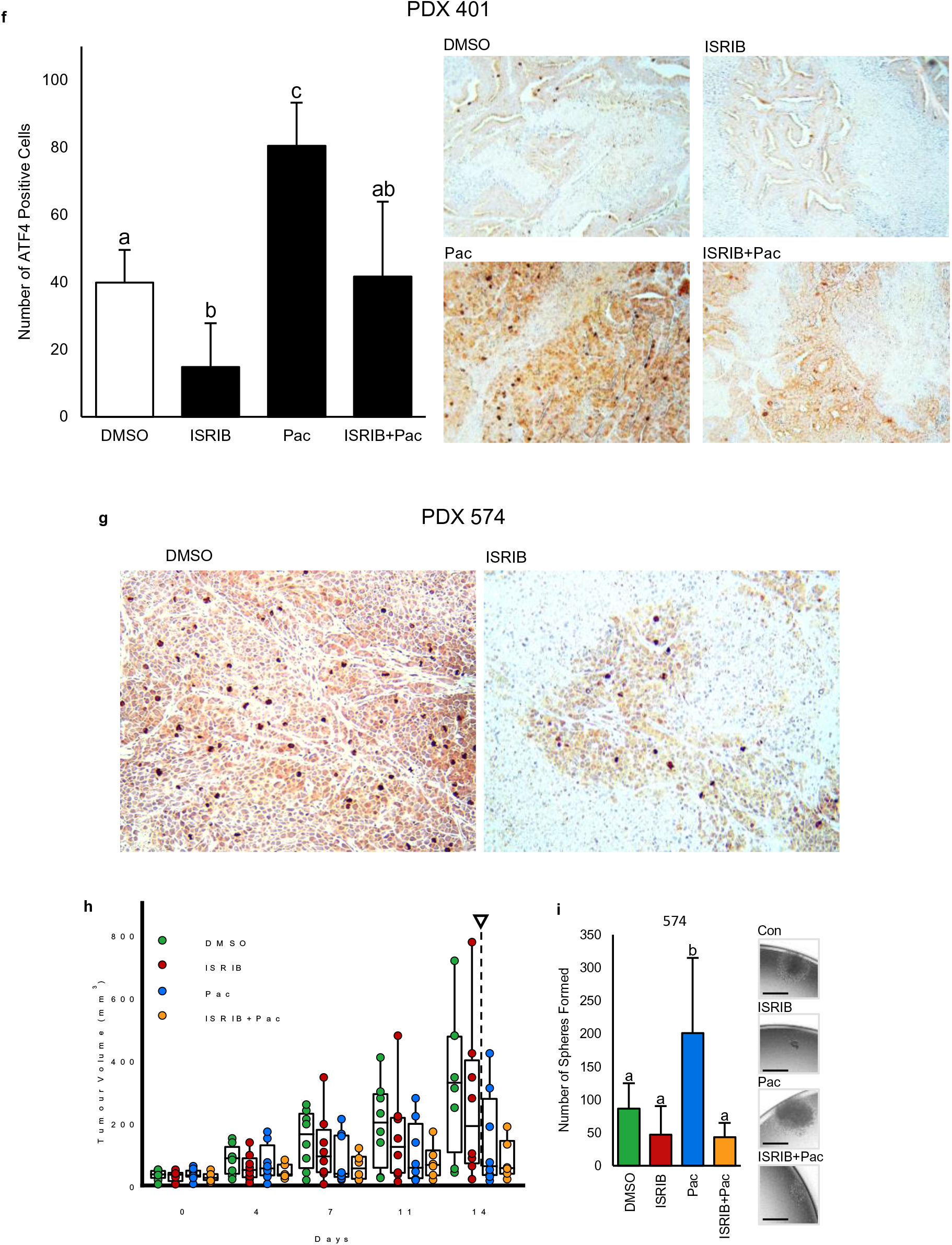
ISRIB mitigates therapy-induced BCSCs and improves efficacy of paclitaxel: **a)** Western blots of eIF2α-p, eIF2α and 4E-BP1 in lysates extracted from SUM149 cells treated for 0-12h with paclitaxel (20nM) show that paclitaxel induces eIF2α phosphorylation and an increase in 4E-BP1. Actin is used as a loading control. **b)** Surviving fractions of SUM149 cells exposed to increasing doses of paclitaxel (5 nM) in the presence or absence of ISRIB (10nM). Bars represent mean surviving fraction ± SD (n=3). Values indicated by an asterisk (*) mark doses wherein ISRIB significantly increased the efficacy of paclitaxel (p<0.05, n=3). **c)** Immunohistochemical detection (brown) of ATF4 in SUM149 tumours from mice treated every second day for 2 weeks with DMSO, ISRIB (2.5 mg/kg), paclitaxel (20 mg/kg), or paclitaxel + ISRIB (2.5 mg/kg). Accompanying bar graph represents the mean number of ATF+ cells per field of view ± SD. Different letters show significant differences (p<0.01, n=7). **d)** Immunohistochemical analysis of CA9 and NODAL (brown) in well and poorly differentiated PDX lines 401 and 574, respectively, showing that PDX 574 is enriched in both CA9 and NODAL. **e)** Histograms of CA9-FITC expression and Nodal APC-A expression of cells dissociated from SUM149 (n=6), PDX401 (n=5) and PDX574 (n=4) tumours. Bar graphs represent differential expression of CA9 in NODAL high and NODAL low populations **f,g)** Immunohistochemical detection (brown) of ATF4 in **f)** 401 and **g)** 574 PDX tumours from mice treated every second day for 2 weeks with DMSO, ISRIB (2.5 mg/kg), paclitaxel (20 mg/kg), or paclitaxel + ISRIB (2.5 mg/kg). Accompanying bar graph represents the mean number of ATF+ cells per field of view ± SD. Letters show significant differences (p<0.01, n=3). For graphs, unles otherwise indicated, values indicated by an asterisk (*) are statistically different from controls and values indicated by letters are statistically different from each other (p<0.05). **h)** PDX574 tumour volumes in mice treated every second day for 2 weeks with DMSO, ISRIB (10 mg/kg oral), paclitaxel (20 mg/kg), or paclitaxel (20 mg/kg) + ISRIB (10 mg/kg oral). Arrows indicate cessation of treatment. Data presented as mean tumour volume ± SD (n=6). **i)** The number of spheres formed from 9600 viable PDX574 cells dissociated from 1cm diameter tumours taken from mice in h) and then cultured in single cell suspensions. Bars represent the mean tumoursphere count ± SD (n=6). Images of representative spheres are presented. Micron bars = 100 μm.

**Supplementary Table 1.**
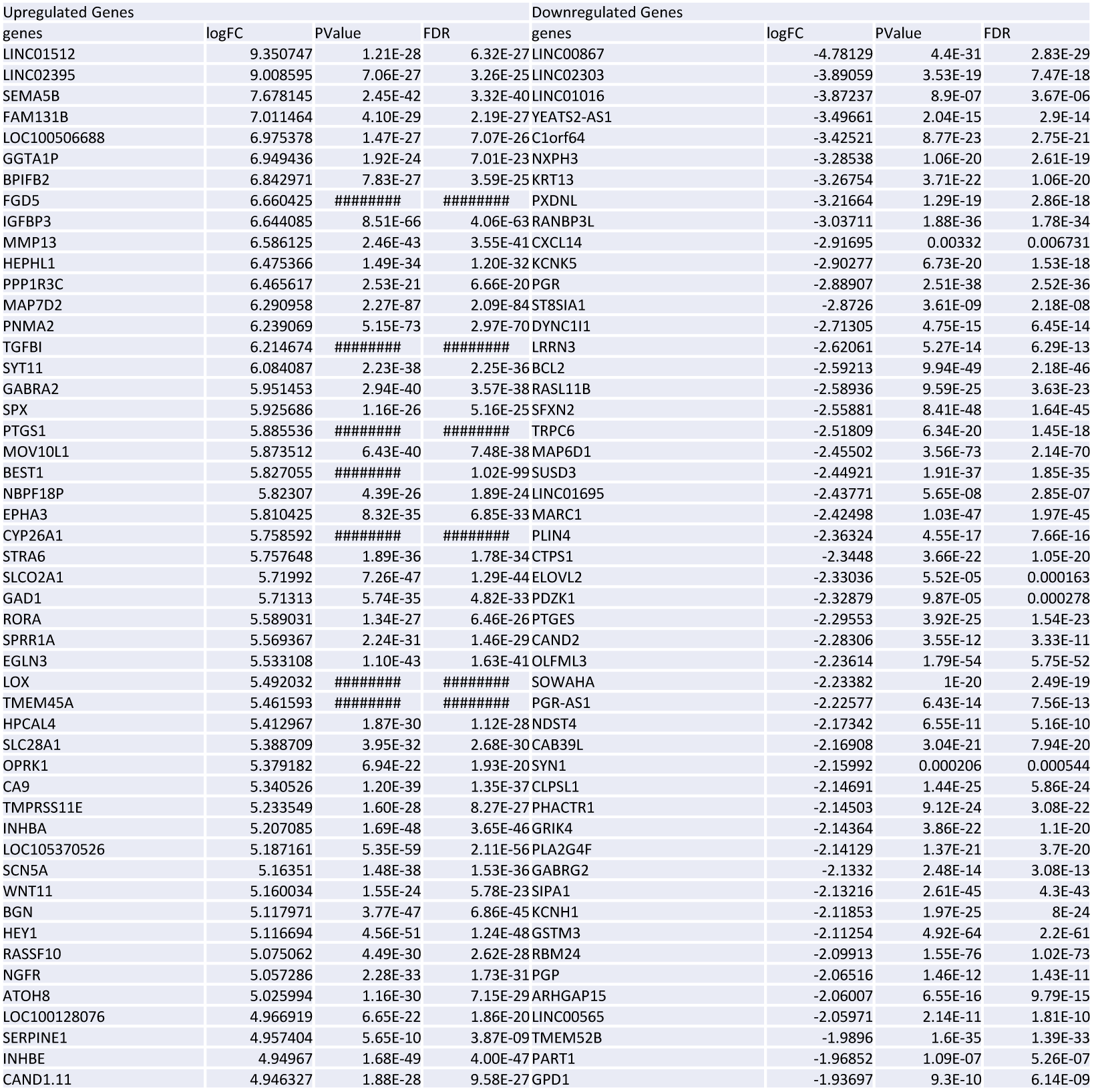
Top 50 Gene Most Altered by Hypoxia in T47D Breast Cancer Cells

**Supplementary Table 2.**
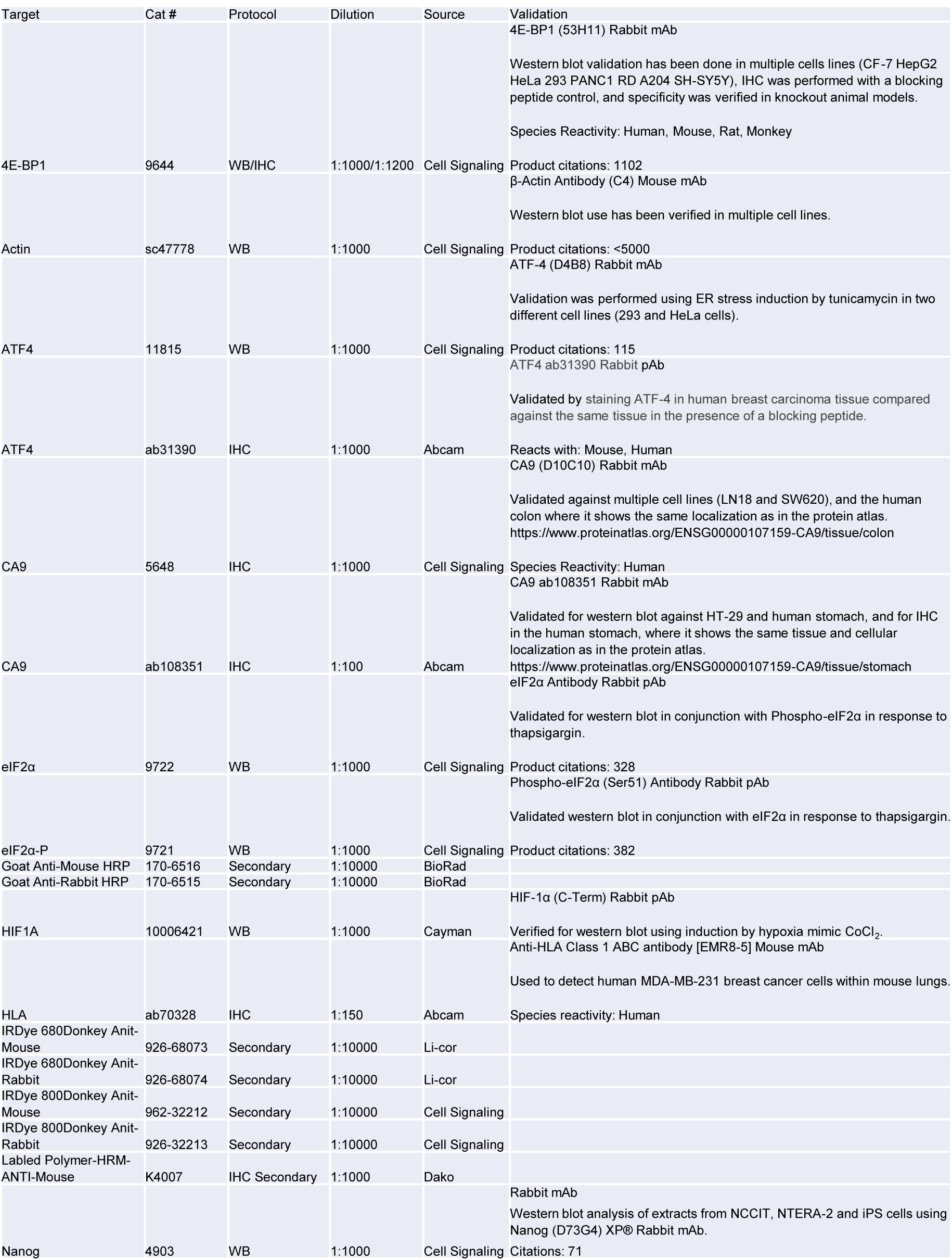
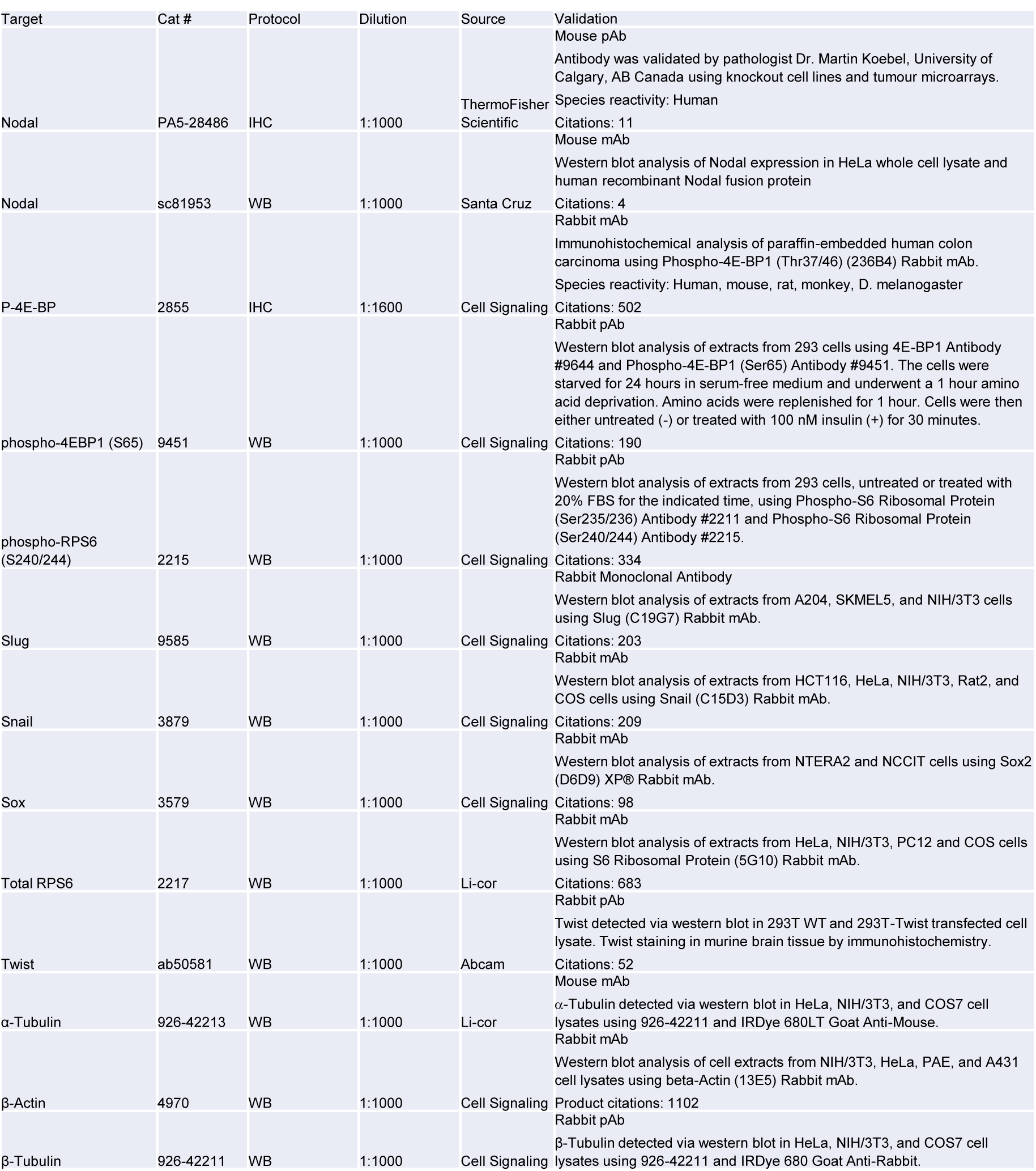
Antibodies Used

**Supplementary Table 3.**
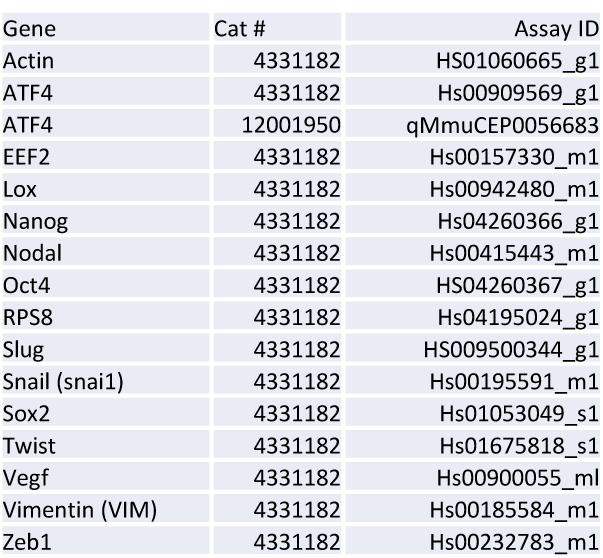
Primer Probes Used

**Supplementary Table 4.**
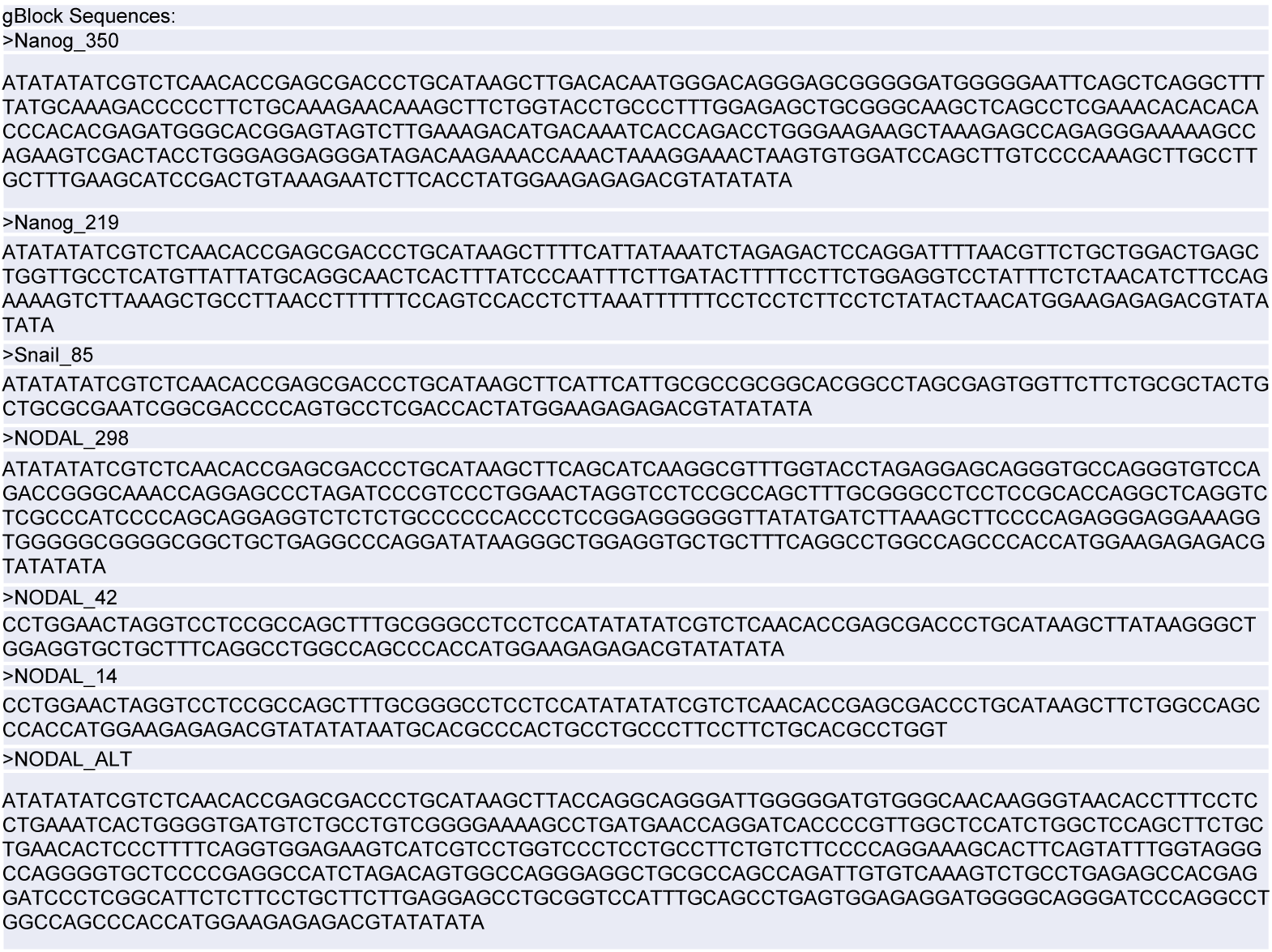
G Blocks Used in Luciferase Reporter Assays

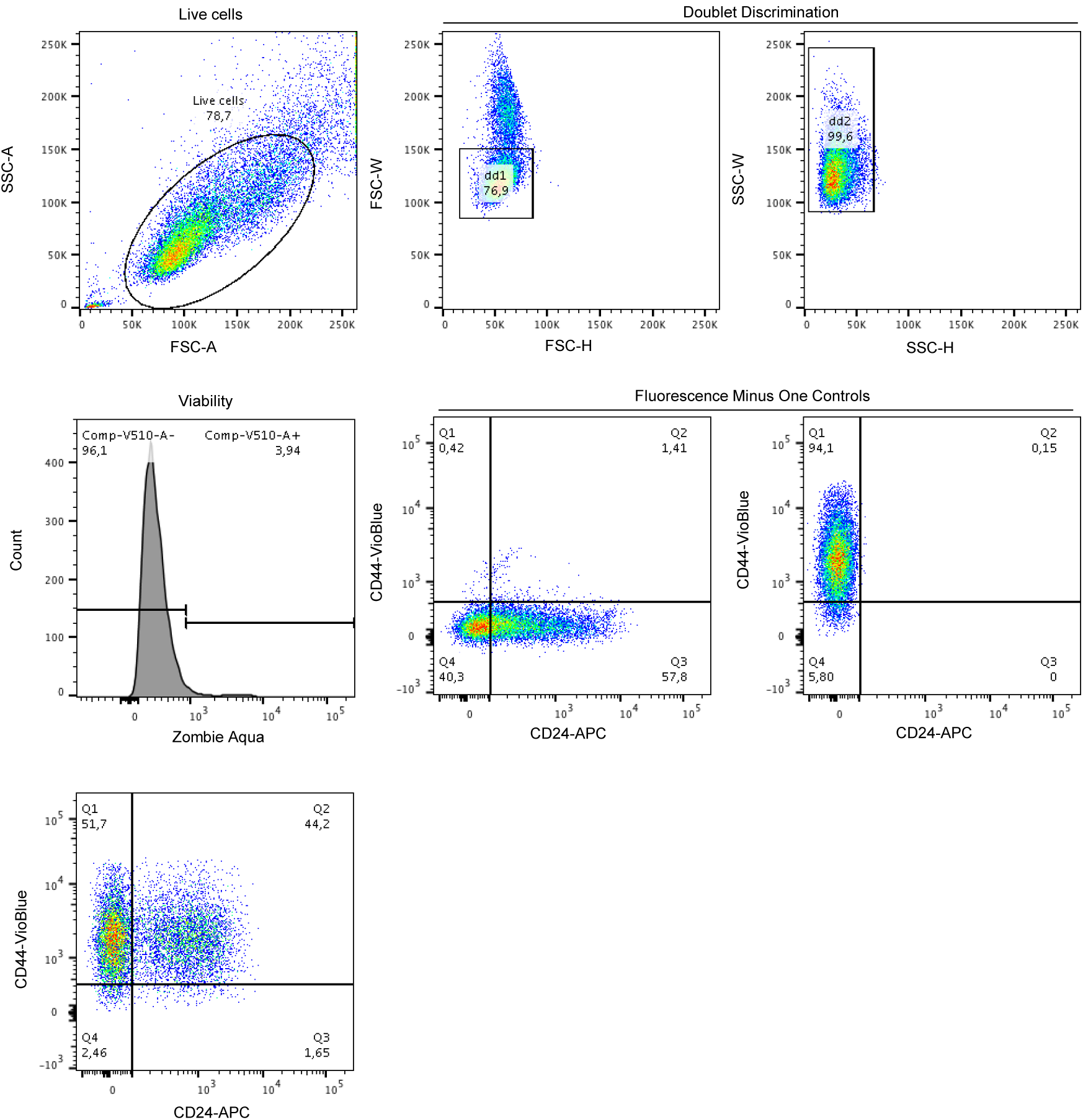
Schema of gating as described in the methods

